# Pathogenic Role of FGFR3 Autoantibodies in Small Fiber Neuropathy

**DOI:** 10.1101/2025.06.01.657230

**Authors:** Lyuba Y Salih, Nicolas LA Dumaire, Clémence Gieré, Erin Vest, Haya Alkhateeb, Christian P Moritz, Yannick Tholance, Jean-Philippe Camdessanché, Jean-Christophe Antoine, Jérôme Honnorat, Jafar Kafaie, Liberty François-Moutal, Aubin Moutal

**Affiliations:** Department of Pharmacology and Physiology, Saint Louis University School of Medicine, Saint Louis, Missouri, USA; Institute for Translational Neuroscience, Saint Louis University School of Medicine, Saint Louis, Missouri, USA; MeLiS | CNRS UMR 5284 | INSERM U1314, Synaptopathies and Autoantibodies, Institut National de la Santé et de la Recherche Médicale (INSERM), U1217, Centre National de la Recherche Scientifique (CNRS) UMR5310, 69000 Lyon, France; Université Claude Bernard Lyon 1, 69000 Lyon, France and University Jean Monnet, 10 rue de Marandière, 42270 Saint-Priest-en-Jarez, France; Department of Neurology, University Hospital of Saint-Etienne, Saint-Etienne, France; Department of Neurology, Saint Louis University School of Medicine, Saint Louis, Missouri, USA

**Author notes:** Correspondence to: Aubin Moutal, PhD, Saint Louis University - School of Medicine, Department of Pharmacology and Physiology, 1402 S. Grand Blvd., Schwitalla Hall, Room 432, Saint Louis, MO 63104, USA.

**Keywords:** FGFR3, autoantibodies, sensory neuron, hyperexcitability, small fiber neuropathy, CRISPR, dorsal root ganglia, neuropathic pain

## Abstract

Sensory neuronopathies (SNN) and small fiber neuropathies (SFN) are debilitating disorders associated with neuropathic pain, yet their underlying mechanisms remain poorly understood. Autoantibodies against fibroblast growth factor receptor 3 (FGFR3-Abs) define a subset of patients with consistent reports of neuropathic pain harboring a distinct clinical phenotype characterized by small-fiber and non-length-dependent neuropathy, suggesting dorsal root ganglia (DRG) dysfunction. FGFR3-Abs bind to sensory neurons within dorsal root ganglia (DRG). The target of autoantibodies FGFR3 is expressed at the transcript and protein level in human sensory neurons, suggesting that FGFR3-Abs could find their target in primary afferents. DRG neurons exposed to FGFR3-Abs rapidly acquired a hyperexcitability phenotype which was linked to mechanical hypersensitivity, mirroring patient-reported pain symptoms. CRISPR mediated gene editing of FGFR3 in sensory neuron prevented FGFR3-Abs induced sensitization of sensory neurons and mechanical hypersensitivity. In parallel, Epitope mapping reveals extracellular FGFR3 epitopes essential for antibody-induced sensitization and pain hypersensitivity. Together this work suggests that beyond their role as biomarkers, FGFR3-Abs are pathogenic in small fiber neuropathy by acting directly on DRG neurons. This positions both FGFR3-Abs and FGFR3 signaling as actionable therapeutic targets for modulating sensory neuron excitability and treating autoimmune painful neuropathies.

## Introduction

Sensory neuronopathies (SNN) and small fiber neuropathies (SFN) are an ensemble of disorders characterized by dysfunctional Aδ fibers, which contribute to mechanosensation and nociception, and unmyelinated C-fibers, which predominantly transmit nociceptive stimuli[1]. The painful symptoms experienced by the patients are often the main contributor to seeking medical intervention and decreased quality of life[2]. Autoantibodies against fibroblast growth factor receptor 3 (FGFR3-Abs) were described as a biological marker for a subgroup of patients with sensory neuronopathy and small-fiber neuropathy[3] with a high prevalence of pain (54%) and paresthesia (80%)[4]. This collection of sensory disorders suggested that seropositivity for FGFR3-Abs may directly contribute to sensory dysfunction and pain sensitization.

FGFR3 is a transmembrane tyrosine kinase receptor expressed in sensory neurons[3a], regulating axonal development and damage-induced sensory neuron survival[5]. FGFR3 activation leads to a wide range of intracellular signaling cascades including p38 mitogen-activated protein kinase (p38), and extracellular signal-regulated kinase (ERK)[6]. Mutations in FGFR3 cause skeletal dysplasias including overgrowth of long bones and vertebrae or several forms of dwarfism, achondroplasia, and Crouzon syndrome[7]. Activating FGFR3 mutations in adults are a common driver of multiple myeloma as well as urothelial and cervical carcinomas[8]. While FGFR3 is known to regulate axonal development and sensory neuron survival, its role in nociception and sensory processing remains unexplored. Given this, we investigated whether these FGFR3 autoantibodies directly modulate DRG neuron excitability and pain responses.

From a retrospective cohort analysis, we observed that FGFR3-Abs patients consistently report neuropathic pain and display a distinct sensory neuropathy phenotype suggesting dorsal root ganglia (DRG) dysfunction. To expand on these clinical findings, we first validated that the target of autoantibodies, FGFR3, is expressed in human sensory neurons. Using FGFR3-Abs seropositive samples from patients with SNN or SFN, we demonstrated that FGFR3-Abs induce sensory neuron hyperexcitability. Furthermore, FGFR3-Abs injection in the paw of rats led to mechanical hypersensitivity, thus showing that human autoantibodies may be the basis of reported pain in patients with SNN or SFN. CRISPR mediated deletion of FGFR3 in DRG neurons demonstrated that the above effects directly engage sensory neurons. Adding to previous studies[9], we mapped the FGFR3 epitopes recognized by autoantibodies using high-density peptide arrays, identifying key extracellular binding sites. This helped us uncover that FGFR3-Abs sensitize sensory neurons via binding to FGFR3 extracellular domain to activate downstream P38 and ERK signaling pathways. Our study demonstrates that FGFR3-Abs bind to and activate FGFR3 in DRG neurons, leading to increased excitability and pain behaviors in rats. Beyond serving as biomarkers for SFN, FGFR3-Abs actively contribute to pain hypersensitivity, making them a potential target for therapeutic intervention.

## Methods

### Materials and Reagents

All chemicals, unless noted were purchased from Sigma (St. Louis, MO). All peptides (>95% purity) were purchased from Genscript (Piscataway, NJ). Extracellular region (ECR) peptide sequences were TGLVPSERVLVGPQR, LNASHEDSGAYSCRQRLTQRV, ERMDKKLLAVPAANTVRF and SWLKNGREFRGEHRIGGIKLR. Juxtamembrane domain (JMD) peptide sequences were SRFPLKRQVSLESNASMSSNTP and LELPADPKWELSRARLTLGKPL. All peptide synthesis was verified by HPLC and mass spectrometry. Peptides were all resuspended in DMSO before use.

### Origin of FGFR3 sera

Patient sera were obtained from the biobank of the University Hospital of Saint-Etienne (CRB42 CHU Saint-Etienne, France, AC 2018-3372, NFS96-900, N° of collection DC-2010-1108). All the sera were stored in aliquots at −80 °C. Detection of FGFR3-Abs was performed by using a previously described indirect ELISA[10].

### Retrospective study of FGFR3 positive patients with SFN

A retrospective chart review was conducted with Institutional Review Board approval on patients diagnosed with painful small fiber neuropathy (SFN) between 2009 and 2017 at Saint Louis University Hospital and Cardinal Glennon Children’s Hospital Neurology Clinics. Inclusion criteria included a clinical SFN diagnosis, robust clinical data, and testing with the Washington University Sensory Neuropathy Panel. Patients with incomplete data or dual positivity for FGFR-3 and TS-HDS antibodies were excluded. Seronegative comparators were consecutive patients from the same timeframe who were negative for all antibodies on the panel (including FGFR3 and TS-HDS) and had complete clinical/biopsy/electrodiagnostic data. No frequency-matching or post-hoc balancing was performed. A total of 172 patients met all criteria and were divided into two cohorts: FGFR-3 positive (n = 63), and seronegative (n = 68). Data collected included demographics, medical history, neuropathic pain and autonomic symptoms, laboratory results, skin biopsy, and EMG/NCS findings. Antibody testing included antibodies against FGFR-3, TS-HDS, and other neural antigens. Data from seronegative and FGFR3 positive patients were analyzed and included in this paper. Data were analyzed using GraphPad Prism 10Analyses included demographics, comorbidities, symptoms, skin biopsy, and lab results. A p-value < 0.05 was considered significant, and results are reported as the frequency of abnormal findings with corresponding p-values.

### Human sera and Dorsal Root Ganglion procurement

Human sera were obtained per standard diagnostic procedures from the University Hospital of Saint-Etienne, France. The respective patients were of similar age, 44 to 61 years old, and without any other confounding autoantibodies, apart from patient #10 who had anti-ganglioside antibodies (**Table S1**). Samples were procured with informed consent of the participants, de-identified, and transferred to the lab at Saint Louis University via overnight Fedex. Research with de-identified patients was performed under an Institutional Review Board exemption #4 at Saint Louis University. Human DRG were obtained within one hour of cross-clamp from Mid America Transplant or Washington University in Saint Louis and transported to Saint Louis University (information about the tissues used in this study can be found in **Table S2**). All studies have been classified as an IRB exemption #4 at Saint Louis University.

### Animals

Adult male and female Sprague-Dawley rats (Pathogen-free, 100–250 g) were purchased from Envigo-Harlan Laboratories (Indianapolis, IN) and housed in environmentally-controlled rooms maintained at 23 ± 3°C with a 12-hour light/dark cycle (food and water available *ad libitum)*. Behavioral experiments were conducted by experimenters blinded to treatment groups. All studies were conducted in accordance with guidelines approved by Saint Louis University in the Guide for the Care and Use of Laboratory Animals (Protocol #: 3014) and the National Institutes of Health.

### gRNA design, cloning and lentiviral packaging for CRISPR/Cas9 targeting FGFR3

Our strategy to delete FGFR3 focused on targeting exon 1 of the rat Fgfr3 gene (Ensembl ID: ENSRNOT00000023144.7) with a guide RNA (gRNA: gCTGCGTGCTAGTGTTCTGCG, on-score 70.5, off-score 44.7) similar to previous studies [11]. The target exon was chosen to achieve complete deletion of the receptor while minimizing off-target Cas9 activity, as previously verified by our group and others. The indicated gRNA sequence was inserted into the Esp3I restriction site of the pL-CRISPR.EFS.GFP plasmid (Addgene plasmid #57819, Cambridge, MA)[12], a construct designed to co-express Cas9, GFP and the gRNA. The control plasmid contained a non-targeting gRNA (gGAGACGTGACCGTCTCT) together with GFP and the Cas9 enzyme. Cloning, plasmid production, lentiviral packaging and quality controls were completed by Genscript.

### Intra-sciatic nerve injections

Intraneural injections was adapted from established procedures in mice [13]. Rats were anesthetized with 5% isoflurane for induction and maintained under 2.5% isoflurane. After confirming areflexia, a ∼2–3 cm incision was made on the right flank, approximately 1 cm lateral to the hip. Sterile wooden toothpicks were used to bluntly dissect the musculature and expose the main branch of the sciatic nerve. To prevent desiccation, a drop of sterile saline was applied to the exposed nerve. A 10 µL Hamilton syringe fitted with a 30 G needle was inserted longitudinally into the sciatic nerve, and 2 µL of lentivirus (10^7^ infectious particles, CRISPR control or CRISPR FGFR3) was injected. The needle was left in place for ∼60 s to allow diffusion before being carefully withdrawn. Muscles were sutured with 4-0 silk, and the skin incision was closed with wound clips. An antibiotic ointment was applied to the wound site. This injection leads to gene editing directly in the DRG [14].

### Testing of mechanical hypersensitivity

The assessment of tactile hypersensitivity (i.e., a decreased threshold to paw withdrawal after probing with normally innocuous mechanical stimuli) consisted of testing the withdrawal threshold of the paw in response to probing with a series of calibrated fine (von Frey) filaments with increasing stiffness, from 0.4 to 15 g. Filaments were applied perpendicularly to the plantar surface of the hind paw and the withdrawal threshold measured by the ‘up-and-down’ method. The withdrawal threshold was determined by sequentially increasing and decreasing the stimulus strength and data were analyzed with the nonparametric method of Dixon, as described by Chaplan et al[15] and expressed as the mean withdrawal threshold.

### Immunohistofluorescence and epifluorescence imaging

Tissues were dissected from adult rats and then fixed using 4% paraformaldehyde for 2h at room temperature (RT). The fixed tissues were transferred into a 30% sucrose solution and left at 4°C until sinking of the tissues could be observed (∼3 days). Tissues were cut at 10 µm thickness on a cryostat and fixed onto charged slides and kept at –20°C until use. Prior to antibody staining, slides were dried at room temperature for 30 min and re-hydrated with phosphate buffered saline (PBS). Next, slices were permeabilized and saturated using PBS containing 3% BSA, 0.1% triton X-100 solution for 30 min at RT, and then sera with FGFR3-Abs were added overnight (1/100) at 4°C. The slices were then washed 3x in PBS, and incubated with PBS containing 3% BSA and secondary antibody (Alexa 555 Goat anti-human IgG - Life Technologies, Cat# A21433) for at least 3h at RT. After 3 washes (PBS, 10 min, RT), slides were mounted and stored at 4°C until analysis. For all experiments, negative controls were added by omission of the primary antibody. This control was used to account for background fluorescence and exposure was set to return no signal in negative controls. Then acquisition parameters were kept constant for all other image acquisitions. Immunofluorescent micrographs were acquired on an EVOS FL Auto (Life Technologies) using a 10x objective. The freeware image analysis program Image J (http://rsb.info.nih.gov/ij/) was used to generate the images after a background removal step.

### Isolation and culture of rat dorsal root ganglion neurons

DRG neurons (all levels) from Sprague-Dawley rats were isolated, excised, and placed in complete DRG medium: Dulbecco Modified Eagle Medium (DMEM, Life technologies) containing penicillin (100 U/mL) and streptomycin (100 µg/mL, Cat# 15140, Life technologies), 30 ng/ml nerve growth factor, and 10% fetal bovine serum (Hyclone, Cat# 25-550). The ganglia were dissociated enzymatically with DMEM (Cat# 11965, Life technologies) solution containing neutral protease (3.125 mg/ml, Cat#LS02104, Worthington) and collagenase Type I (5 mg/ml, Cat# LS004194, Worthington) in a 37°C, 5% CO_2_ incubator for 30 min. The dissociated cells were resuspended in complete DRG medium. Cells were then seeded on poly-D-lysine (Cat# P6407, Sigma) coated glass coverslips (Cat# 72196-15, electron microscopy sciences) and placed in a 37°C, 5% CO2 incubator for 45–60 min. Cultured cells were then flooded gently with complete DRG medium. Treated DRG neurons were incubated with FGFR3-Abs (1:100), FGFR3 interacting peptides (100 ng/ml) or corresponding vehicles for 30 minutes as indicated.

### Calcium imaging

DRG neurons were loaded at 37°C with 3 µM Fura-2AM (Cat#F-1221, Life technologies, stock solution prepared at 1 mM in DMSO, 0.02% pluronic acid-127, Cat#P-3000MP, Life technologies) for 30 minutes (λ_ex_ 340, 380 nm/λ_em_ 512 nm) to follow changes in intracellular calcium ([Ca^2+^]) in Tyrode’s solution (at ∼310 mOsm) containing 119 mM NaCl, 2.5 mM KCl, 2 mM MgCl_2_, 2mM CaCl_2_, 25 mM HEPES, and 30 mM glucose (pH 7.4). Baseline was acquired for 1 minute followed by stimulation (10 sec) with stimulatory solutions including; 100 nM Capsaicin, 40 mM KCl, 10 µM ATP, and 170 mOsm trigger (52 mM NaCl, 2.5 mM KCl, 2 mM CaCl_2_, 25 mM HEPES, and 30 mM glucose, pH 7.4). Fluorescence imaging was performed with an inverted microscope, Nikon Eclipse TE2000 (Nikon Instruments Inc.), using objective Nikon Super Fluor MTB FLUOR 4× 0.50 and a Nikon Camera (Nikon Instruments Inc.) controlled by NIS Elements software (version 4.20, Nikon instruments). The excitation light was delivered by a Lambda-421 system (Sutter Instruments). The excitation filters (340±5 nm and 380±7 nm) were controlled by a Lambda 10-2 optical filter change (Sutter Instruments). Fluorescence was recorded through a 505 nm dichroic mirror at 535±25 nm. The changes in [Ca^2+^] were monitored by following the ratio of F340/F380, calculated after subtracting the background from both channels.

### Human DRG preparation and immunostaining

Human (T12-L2) DRG were obtained from Mid America Transplant and supplied to the Moutal laboratory at St. Louis University as before[16]. All studies involving human tissues have been classified under IRB exemption #4 at Saint Louis University. The tissues were fixed by immersion in 10% formalin for at least one day. After fixation, the DRGs were trimmed of their connective tissue and roots, cut in half, prior to embedding into paraffin blocks. The paraffin blocks were sectioned in 5 µm slices. Deparaffination is carried out by three xylene washes for 5 minutes each, followed by two 100% ethanol washes for 1 minute each, then two 95% ethanol washes for 1 minute each, and finally by washing in running water for 2 minutes.

For antibody staining, the slides were washed twice with PBS for 5 minutes each at RT, then exposed for 30 minutes to a blocking solution containing 5% donkey serum and 1% BSA in PBS at room temperature. The primary antibody against FGFR3 (Sigma, Cat# SAB5702794, 1:100) was then incubated in blocking solution, overnight at 4°C in a humidified chamber. The slides were then washed 3 times, 15 minutes each with PBS at room temperature before adding the secondary antibody solution (donkey anti-rabbit, IgG, Alexa Fluor® 594, 1/300) for one hour at room temperature. Slides were then washed with PBS 3 times for 15 minutes at room temperature. Finally, slides were mounted in ProLong Gold (Thermo Fisher Scientific, #P36930) to protect them from fading and photobleaching. A no-primary-antibody control was done as described above. Immunofluorescent micrographs were acquired on a Leica SP8 inverted upright microscope using a 10x dry objective. For a clearer and more reliable signal, the acquisition window of the detector was set between 602 and 637 nm, which corresponds to 15 nm above and below the emission peak of Alexa 594. For all quantitative comparisons among cells under differing experimental conditions, camera gain and other relevant settings were kept constant. The freeware image analysis program Image J (http://rsb.info.nih.gov/ij/) was used for contrast enhancement and extracting representative pictures.

### Nerve terminal enrichment and fractionation from rat skin

Nerve terminals in skin tissues are biochemically similar to synaptic terminals. As such they can be isolated using well defined existing protocols for synaptosome fractionation [17]. Adult rats were sacrificed by isofluorane overdose and decapitation, the glabrous skin from the hindpaws dissected. Fresh tissues were homogenized using a Dounce homogenizer in ice-cold Sucrose 0.32M, HEPES 10 mM, pH 7.4 buffer. The homogenates were centrifuged at 1000xg for 10 min at 4°C to pellet the insoluble material. The supernatant was harvested and centrifuged at 12000xg for 20 min at 4°C to pellet a crude membrane fraction. The pellet was then re-suspended in a hypotonic buffer (4 mM HEPES, 1 mM EDTA, pH 7.4) and the resulting synaptosomes pelleted by centrifugation at 12000xg for 20 min at 4°C. Nerve terminal enrichment was verified by immunoblotting for PSD95, TRPV1 and NaV1.8 as markers of synaptic terminals, C-fibers and nociceptors respectively (**Figure 3A**). All buffers were supplemented with protease and phosphatase inhibitor cocktails. Protein concentrations were determined using the BCA protein assay.

### Immunoblotting of rat and human dorsal root ganglia or skin fractions

Rat DRG were dissected and proteins extracted by sonication in 50 mM Tris-HCl, pH 7.4, 50 mM NaCl, 2 mM MgCl_2_, 1% [vol/vol] NP40, 0.5% [mass/vol] sodium deoxycholate, 0.1% [mass/vol] SDS with Protease inhibitors (Cat# B14002; Selleck Chemicals, Houston, TX), phosphatase inhibitors (Cat# B15002, Selleck Chemicals), and Pierce™ Universal Nuclease (Cat# 88701, Thermo Fisher Scientific). In the case of human DRG, the same lysis buffer was used and samples were first homogenized then sonicated. Lysates were clarified at 10,000xg, 4°C, 10min and protein concentrations determined using the BCA protein assay (Cat# PI23225, Thermo Fisher Scientific, Waltham, MA). Indicated samples (20 µg of total protein each) were loaded on 4-20% Novex® tris-glycine gels (Cat# XP04205BOX, Thermo Fisher Scientific, Waltham, MA). Proteins were transferred for 1h at 100 V using tris glycine sodium docecyl sulfate buffer (TGS, 25 mM Tris pH=8.5, 192 mM glycine, 0.1% (mass/vol) SDS), 20% (vol/vol) methanol as transfer buffer to polyvinylidene difluoride (PVDF) membranes 0.45μm (Cat# IPVH00010, Millipore, Billerica, MA), pre-activated in pure methanol. After transfer, the membranes were blocked at room temperature for 1 h with TBST (50 mM Tris-HCl, pH 7.4, 150 mM NaCl, 0.1

% Tween 20), 5% (mass/vol) non-fat dry milk, then incubated separately in the primary antibodies FGFR3 (Sigma, Cat#SAB5702794), p38 (Cell Signaling, Cat#8690), pp38 T180/Y182 (Cell Signaling, Cat#4511), JNK1 (Cell Signaling, Cat#3708), pJNK1 T183/Y185 (Cell Signaling, Cat#4668), ERK (Cell Signaling, Cat#4696), pERK T202/Y204 (Cell Signaling, Cat#4370), PSD95 (Thermo Fisher Scientific, Cat#MA1-045), NaV1.8 (Alomone, Cat#ASC-016), TRPV1 (Thermo Fisher Scientific, Cat#PA1-29770), Actin (Thermo Fisher Scientific, Cat#PA116889) or βIII-Tubulin (Promega, Cat#G7121) in TBST, 5% (mass/vol) BSA, overnight at 4°C. Following incubation in horseradish peroxidase-conjugated secondary antibodies from Jackson immunoresearch, IgG, IgA, IgM, blots were revealed by enhanced luminescence (WBKLS0500, Millipore, Billerica, MA) before exposure to photographic film. Films were scanned, digitized, and quantified using Un-Scan-It gel version 7.1 scanning software by Silk Scientific Inc.

### RNAscope in situ hybridization

Immediately following procurement, human DRG tissues were immediately added to 10% formalin for fixation. Samples were then trimmed of connective tissue, processed into paraffin, and made into tissue microarrays (TMA, 7 donors per slide) using a 5 mm punch and sectioned into 4.5 µm slices. Following fixation and TMA selection, slices were then stained according to RNAscope™ Multiplex Fluorescent Detection Kit v2 (Cat# 323110, Advanced Cell Diagnostics, Newark, CA), using the following RNAscope probes; Hs-FGFR3 (Cat# 310791), and Hs-TRPV1 (Cat#415381). TMAs were then incubated with DAPI (10 µg/ml) for 10 min, washed with 1X PBS and air dried prior to mounting with c mounting medium (Thermo Fisher, Cat# P36934). In our experiments, we included RNAscope positive and negative controls per manufacturer’s instructions. Human tissue TMAs were imaged using Leica DM6 with 40X magnification. We verified the integrity of our staining by positive signals in our positive control and the absence of RNAscope puncta in our negative control. Only clearly identifiable DRG neurons with intact DAPI-stained nuclei and boundaries were included. FGFR3 puncta were scored as small, discrete signals within neuronal somata, while lipofuscin was excluded based on its large size and autofluorescence across channels. To quantify the number of signals for each RNAscope probe within the human DRG TMAs, images were uploaded and thresholded to show only the RNAscope signals in the Omero OME (Open Microscopy Environment) online software. FGFR3 puncta were then counted manually.

### Peptide SPOT array synthesis and hybridization

As previously described[18], peptide spot arrays spanning the sequence of FGFR3 (UniProt ID: P22607, PDBID: 3GRW^18^) were synthesized in 15-mers with a three amino acid increments using the SPOTS-synthesis method. Standard 9-fluorenylmethoxyl carbonyl (Fmoc) chemistry was used to synthesis the peptides and spot the peptides onto Celluspots nitrocellulose membranes that were prederivatized with a polyethylene glycerol spacer (Intavis). Following peptide synthesis, the side chains were deprotected using 80% trifluoroacetic acid (TFA), 3% triisopropylsilane (TIPS), 12% dichloromethane (DCM), 5% H_2_O for 2 hours and then solubilized in in 88.5% TFA, 4% trifluoromethanesulfonic acid (TFMSA), 2.5% TIPS, and 5% H_2_O overnight at room temperature. Peptides were then precipitated with tert-butyl-methyl ether in a 4:1 ratio. The cellulose-peptide conjugates were centrifuged, the supernatant removed, and pellets left to dry before being redissolved in 100% DMSO. The resulting peptides were spotted on membranes fitted to glass microscope slides by the Intavis Multipep robot.

### Immunoblotting peptide arrays

Celluspots slides were washed in TBST (50 mM Tris-HCl, pH 7.4, 150 mM NaCl, 0.1 % Tween 20) for 10 min and blocked in TBST containing 5% (w/v) nonfat dry milk for 1h at room temperature. Patient sera were diluted 1:100 in TBST, 5% nonfat dry milk and incubated on the Celluspot slide overnight. The array was washed 3 times in TBST for 5 min at room temperature and incubated with the Peroxidase AffiniPure™ Goat Anti-Human (Cat#109-035-064, IgG, IgM, IgA) diluted at 1:1000 in TBST, 5% BSA for 2h at room temperature for visualization. The arrays were then washed 3 times each for 5 min in TBST and finally visualized by infrared fluorescence. All arrays contained duplicates that were averaged and then normalized on the maximal value array for each patient. Positive reactivity of the autoantibodies was mapped onto FGFR3 structure (PDBID: 3GRW^18^) to visualize their binding distributions using PyMOL 3.0.2.

### Immunostaining on cultured rat sensory neurons

Immunocytofluorescence was performed on cultured rat DRG neurons. Briefly, cells were fixed using 4% paraformaldehyde for 10 minutes at room temperature. Permeabilization was done as indicated by a 30-minute incubation in PBS, 0.1% Triton X-100, 3% BSA. Cell staining was performed with human sera with FGFR3-Abs (1/100) in PBS with 3% BSA overnight at 4°C. Cells were then washed three times in PBS and incubated with PBS containing 3% BSA and secondary antibodies (Alexa 488 goat anti-human IgG, Cat# A11013 from Life Technologies) for 1 hour at room temperature. Nuclei were stained with DAPI (10 µg/ml) for 10 min and slides mounted in Prolong Diamond (Life technology, Cat# P36961). Images were acquired on a Leica DM6 with 40X magnification with light intensity, gain, exposure and relevant settings kept constant. Analysis was done with image J to quantify the mean fluorescence intensity on raw pictures. Representative pictures were generated with Image J.

### Patch Clamp Electrophysiology

Rat DRG neurons were cultured as described above and spotted onto 12 mm poly-D-lysine coated glass coverslips. Whole-cell patch clamp recordings were performed using an EPC 10 Amplifier-HEKA (HEKA Elektronik, Ludwigshafen, Germany) linked to a computer with Patchmaster software. Extracellular recording solution consisted of: 145 mM NaCl, 3 mM KCl, 2 mM CaCl_2_, 1 mM MgCl_2_, 10 mM HEPES, 10 mM Glucose, pH 7.4 (adjusted with KOH, 310 mOsm). The intracellular recording solution consisted of the following: 130 mM K-Gluconate, 5 mM KCl, 5 mM NaCl, 3 mM Mg-ATP, 10 mM HEPES, 0.3 mM EGTA, 10 mM Glucose, pH 7.3 (adjusted with KOH, 290 mOsm). Action potentials were evoked by injecting depolarizing pulses (1000 ms) from −10 pA to 190 pA in 10 pA increments. The rheobase or minimum current to fire a single action potential, was determined by injecting short depolarizing pulses (50 ms) starting at −10 pA in 10 pA increments until a single action potential was recorded. Capacitive artifacts were fully compensated, and series resistance was compensated by ∼60%, filtered at 30 kHz and digitized at 20 kHz. Fire-polished recording pipettes were pulled from 1.5 mm borosilicate glass capillaries from a horizontal puller (Sutter Instruments, Model P-97) to a resistance of 2 to 6 MΩ. Only small cells with a capacitance of less than 30 pF were considered as C-fibers and included in analysis. The DRG neurons with a resting potential more hyperpolarized than −40 mV, stable baseline recordings, and evoked spikes that overshot 0 mV were used for experiments and analysis. All experiments were performed at room temperature.

### Statistics

Graphing and analysis were completed in GraphPad Prism (Version 10). Details of each analysis are included within the respective figure legends and in **Table S3**. All data sets were checked for normality using D’Agostino & Pearson test. All data plotted represent mean ± SEM. Table 1 data were analyzed using χ² tests for categorical variables. For electrophysiological recordings, data was compared using Mann-Whitney tests (rheobase), One-way ANOVA with the Tukey post hoc test and Kruskal–Wallis test with Dunnett’s post hoc comparisons; for sensory neuron excitability, statistical differences between groups were determined using multiple Mann-Whitney test and Mann-Whitney test. Statistical significance of hypersensitivity was compared by Kruskal-Wallis test followed by the Dunn post hoc test. Behavioral data with a time course were analyzed by two-way ANOVA.

**Table 1:** FGFR3 Seropositive vs Seronegative small fiber neuropathy. Demographics, clinical features, and test findings for FGFR3 seropositive small fiber neuropathy (SFN) patients compared to seronegative SFN patients (control group). All values are frequencies (percent of group) unless otherwise noted. Statistically significant differences (P < 0.05) between the FGFR3 group and controls are indicated. IBS: irritable bowel syndrome, SNAP: sensory nerve action potential, ESR: Erythrocyte Sedimentation Rate.

| Variable | FGFR3+ SFN<br>(n = 63) | Seronegative SFN<br>(n = 68) | OR (95% CI) | p-value | Test |
| --- | --- | --- | --- | --- | --- |
| Age, mean ± SD<br>(years) | 52.1 ± 16.7 | 46.2 ± 13.0 | — | 0.0267 | Welch's t-test (two-<br>sided) |
| Female sex | 42/63 (66.7%) | 58/68 (85.3%) | 0.34 (0.15–<br>0.81) | 0.0122 | Pearson $\chi^2$ |
| Diabetes mellitus | 11/63 (17.5%) | 10/68 (14.7%) | 1.23 (0.48–<br>3.12) | 0.668 | Pearson $\chi^2$ |
| Sjögren's<br>syndrome | 1/63 (1.6%) | 3/68 (4.4%) | 0.35 (0.04–<br>3.45) | 0.620 | Fisher's<br>exact |
| IBS | 1/63 (1.6%) | 10/68 (14.7%) | 0.09 (0.01–<br>0.75) | 0.0068 | Pearson $\chi^2$ |
| Anxiety or<br>Depression | 27/63 (42.9%) | 34/68 (50.0%) | 0.75 (0.38–<br>1.49) | 0.413 | Pearson $\chi^2$ |
| Rheumatoid<br>arthritis | 5/63 (7.9%) | 10/68 (14.7%) | 0.50 (0.16–<br>1.55) | 0.224 | Pearson $\chi^2$ |
| Abnormal sural<br>SNAP | 33/63 (52.4%) | 10/68 (14.7%) | 6.38 (2.77–<br>14.68) | 4.47$\times 10^{-6}$ | Pearson $\chi^2$ |
| Abnormal radial<br>SNAP | 10/63 (15.9%) | 2/68 (2.9%) | 6.23 (1.31–<br>29.65) | 0.0104 | Pearson $\chi^2$ |
| Non-length-dependent SFN<br>(NLDSFN) | 35/63 (55.6%) | 12/68 (17.6%) | 5.83 (2.63–<br>12.95) | 8.84$\times 10^{-6}$ | Fisher's<br>exact |
| Length-dependent<br>SFN (biopsy) | 28/63 (44.4%) | 56/68 (82.4%) | 0.17 (0.08–<br>0.38) | 8.84$\times 10^{-6}$ | Fisher's<br>exact |
| Dysautonomia | 36/63 (57.1%) | 53/68 (77.9%) | 0.38 (0.18–<br>0.81) | 0.0110 | Pearson $\chi^2$ |
| Fibromyalgia | 9/63 (14.3%) | 27/68 (39.7%) | 0.25 (0.11–<br>0.60) | 0.0010 | Pearson $\chi^2$ |
| ESR elevated | 10/63 (15.9%) | 17/68 (25.0%) | 0.57 (0.24–<br>1.35) | 0.197 | Pearson $\chi^2$ |

## Results

### FGFR3 Seropositive Neuropathy Reveals A Distinct Clinical Profile Typical Of DRG Neuron Dysfunction

To establish whether FGFR3-Abs contribute to sensory neuron pathology, we first analyzed clinical characteristics of patients seropositive for FGFR3. Compared to a control cohort of seronegative small fiber neuropathy (SFN) patients, FGFR3-positive individuals exhibited a distinct phenotype marked by fewer systemic comorbidities and an atypical sensory neuropathy profile (**Table 1**). All FGFR3-Abs patients reported neuropathic pain as a primary symptom, consistent with the high prevalence of pain described in previous studies of this patient population. These symptoms were often the main motivator for clinical evaluation and diagnostic testing. FGFR3-positive patients were slightly older on average (52.1 ± 16.7 vs 46.2 ± 13.0 years) and more likely to be male (33.3% vs 15%, *P* = 0.02) (**Table 1**). They had lower rates of fibromyalgia, anxiety/depression, IBS, and autoimmune conditions such as Sjögren’s syndrome and rheumatoid arthritis, suggesting a narrower disease spectrum with less systemic inflammation. Despite this, FGFR3-Abs patients exhibited objective evidence of more pronounced sensory dysfunction (**Table 1**). Specifically, 52.7% had abnormal sural sensory nerve action potentials (SNAPs), and 15.6% showed radial SNAP abnormalities—both significantly elevated compared to seronegative controls (*P* < 0.0001 and *P* = 0.0006, respectively). Additionally, only 45.2% displayed length-dependent small fiber loss on skin biopsy, a hallmark of classical SFN, compared to 63% of controls (*P* = 0.0386) (**Table 1**). This divergence is consistent with a non-length-dependent process which was prominent in FGFR3-Abs SFN (55.6%) for 17.6% of seronegative SFN and raises the possibility of neuronal damage occurring at the level of the dorsal root ganglia (DRG) rather than through distal axonopathy. Taken together, these clinical observations support the idea that FGFR3-positive neuropathy constitutes a distinct clinical entity characterized by consistent pain symptoms, reduced systemic immune features, less distal axonal degeneration, and prominent small-fiber and proximal involvement. Based on these clinical observations, we hypothesized that FGFR3 autoantibodies may act directly on sensory neurons in the DRG to drive pain hypersensitivity.

### FGFR3 Autoantibodies Can Find Their Target In DRG Neurons

To determine whether FGFR3-Abs could act on DRG sensory neurons, we first validated that they can find their autoantigen in rat tissues. Similar to what was described before[3a], FGFR3-Abs from different patients (**Table S1**) all recognized neuronal somata of rat DRG neurons (**Figure 1A**). The immunoreactivity was restricted to sensory neurons and fibers with no signal detected in satellite glia. We next immunostained human DRG (**Table S2**) with a commercial antibody against FGFR3 (**Figure S1**). We found consistent positive signal in neuronal cell bodies across multiple donors, regardless of age or sex (**Figure 1B**). Again, the signal was localized predominantly to neuronal somata and axonal projections with notable staining of surrounding glia. Finally, FGFR3 protein expression in human DRG was validated with the same commercial antibody against FGFR3 (**Figure 1C**). No difference of expression was detected between male and females in our dataset (**Figure 1D**). These observations indicate that FGFR3 is consistently expressed in sensory neurons in both rat and human. Therefore, human FGFR3-Abs could find their target in human sensory neurons which might underpin the increased incidence and intensity of pain in this patient population.

**Figure 1:**
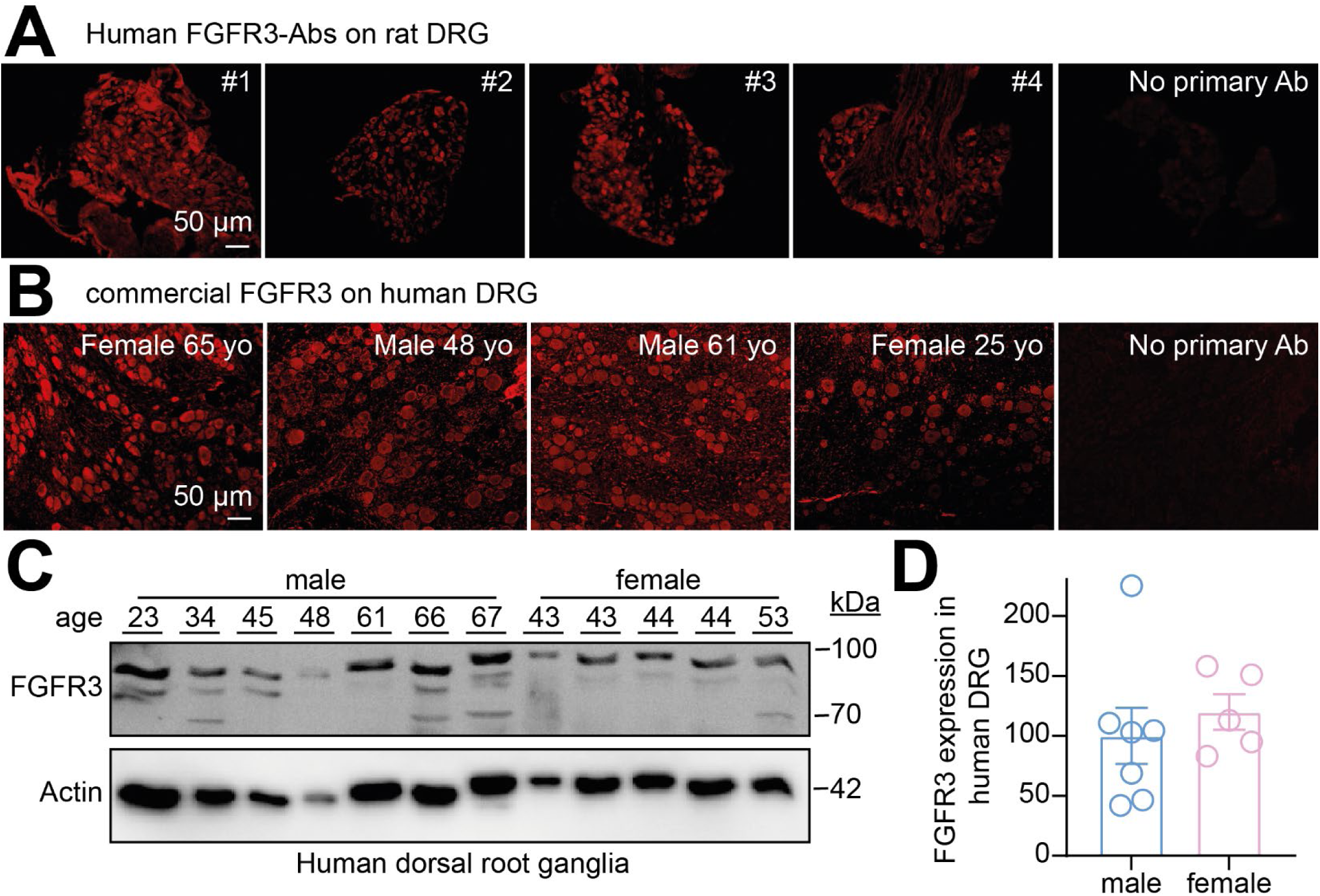
FGFR3-Abs staining and FGFR3 expression in rat and human dorsal root ganglia. (**A**) Representative fluorescent micrographs of rat dorsal root ganglia (DRG) immunolabeled with FGFR3-Abs from the indicated patient (from **Table S1**) showing positive staining within the soma. Patient # is indicated on the top right corner. Scale bar is 50µm. Omission of the primary antibody (Ab) was used as a negative control. (**B**) Representative micrographs of FGFR3 immunostaining with the use of commercial antibody on human DRG tissues from 4 different donors (**Table S2**) confirming the presence of FGFR3 protein within human sensory neurons. Basic donor information is indicated in the top right corner. Scale bar is 50µm. Omission of the primary antibody (Ab) was used as a negative control. (**C**) Immunoblots showing FGFR3 expression in human DRG from 12 donors (7 males, 5 females). Actin was used as a loading control. (**D**) Bar graph with scatter plot of the relative quantification of FGFR3 expression normalized to actin as a loading control in human DRG from male or female donors as indicated. No difference was observed. Mean ± SEM, Mann-Whitney test.

### FGFR3 Expression In Human Nociceptive Neurons

Given the established presence of FGFR3 in DRG, we next sought to determine its distribution among sensory neuron subtypes. First, we mined publicly available data from a spatial transcriptomic study on human DRG[19] which showed the presence of FGFR3 transcripts in several populations of nociceptors and Aδ fibers (**Figure S2**). We next validated this observation by directly assessing FGFR3 expression in DRG neurons including those expressing transient receptor potential vanilloid 1 (TRPV1) as a marker of nociceptive neurons[20]. For this experiment, we used RNAscope to validate the expression of transcripts and provide a secondary validation of FGFR3 gene expression in sensory neurons beside antibody staining and high-throughput RNA-seq (**Figures 2A and S2**). We obtained 28 human DRG (**Table S2**). In all tissues, we found positive staining for FGFR3 in all sensory neurons including nociceptors labeled with TRPV1 (**Figure 2A**). Lipofuscin staining was also prevalent but could be easily identified as marked by asterisks and excluded from quantification (**Figure 2A**). Appraisal of FGFR3-positive neuron size distributions showed that these cells were predominantly small-to medium-diameter neurons (**Figures 2B–D**). This pattern was consistent across both male and female donors, with no significant differences between sexes. We also observed some RNAscope signal for FGFR3 outside sensory neurons which could indicate a level of transcript expression in satellite glia or other circulating cells, which was not detected by our immunostaining (**Figure 1**). These results confirm that FGFR3 is expressed in sensory neurons.

**Figure 2.**
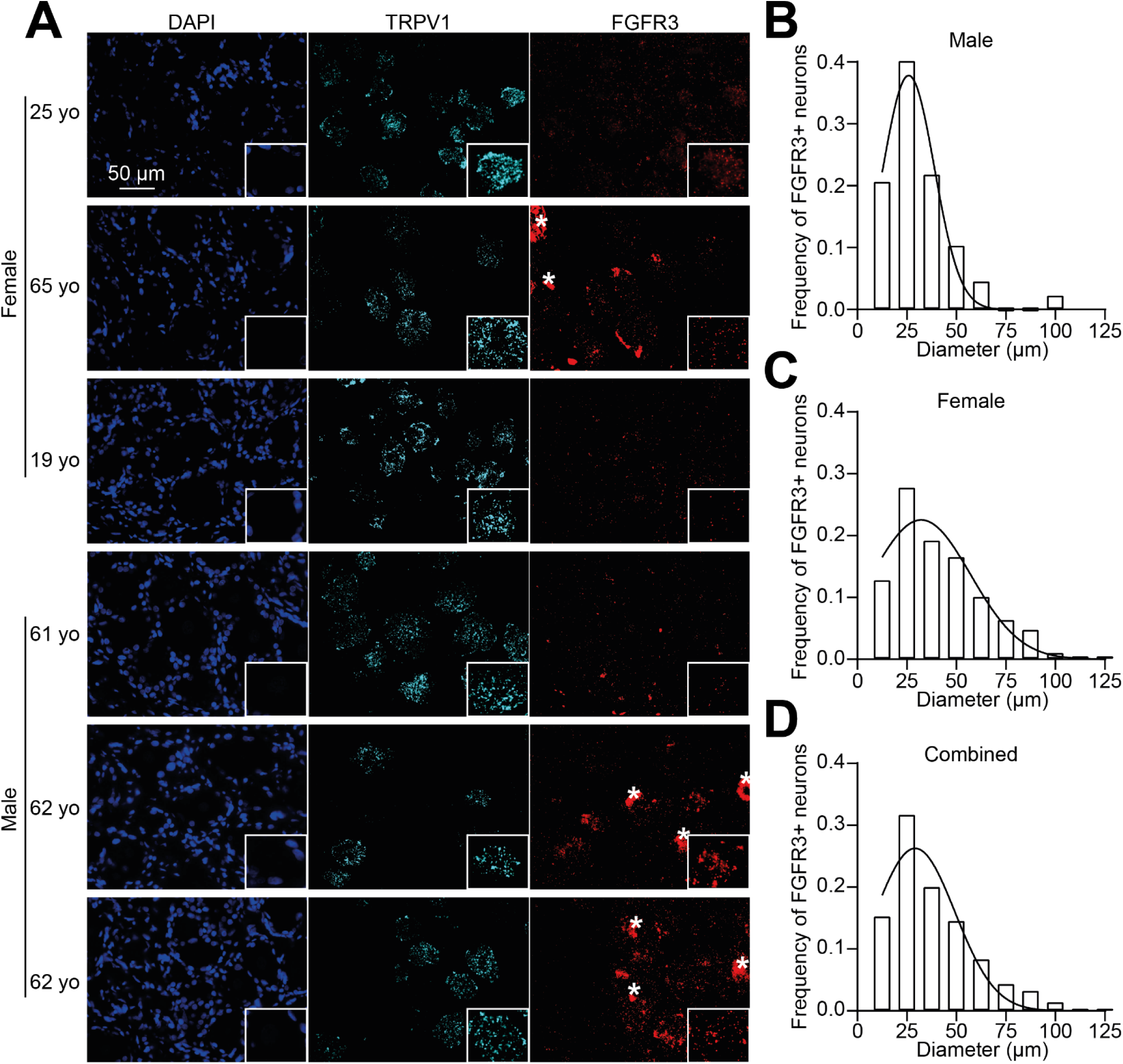
FGFR3 transcripts are expressed within human sensory neurons. (**A**) Representative RNAscope in situ hybridization images of dorsal root ganglia (DRG) revealing TRPV1 (maker of nociceptors, cyan) and FGFR3 (red) transcripts in male and female donors as indicated. DAPI (blue) was used to stain nuclei. Asterisks (*) show lipofuscin signal that was excluded from analysis. Scale bar is 50 µm. Insets show magnified views of TRPV1 and FGFR3 co-expressing neurons in each tissue. Bar graph representing the size distribution of human DRG neurons positive for FGFR3 in (**B**) male or (**C**) female tissues and (**D**) across all donors (combined) showing a prevalence in small-to medium-sized neurons.

### FGFR3 Autoantibodies Induce Mechanical Hypersensitivity

Because FGFR3-Abs are highly correlated with an increased prevalence of pain in patients, we wanted to determine whether FGFR3-Abs could sensitize pain behaviors in rats. To directly reach the sensory nerve terminals in the skin where reflexive testing is performed, we opted for an intraplantar injection of a 1/10 dilution of FGFR3-Abs patient sera as we did before [21]. First, we verified that FGFR3 was expressed on the nerve terminals present in the glabrous skin of the hindpaw. We used principles of synaptosome fractionation and successfully enriched NaV1.8 and TRPV1 as markers of nociceptive nerve terminals (**Figure 3A**). FGFR3 co-fractionated with our markers including PSD95 as an integral component of nerve terminals (**Figure 3A**). Injection of FGFR3-Abs-positive sera from four different patients consistently led to a significant and sustained reduction in paw withdrawal thresholds compared to saline and serum from a healthy control obtained from the blood bank (**Figure 3B**). This mechanical hypersensitivity peaked at three hours after injection and lasted for at least 24 hours and up to 72 hours for two out of four patient samples (**Figure 3B**). Integration of the curve relative to the pre-injection baseline further confirmed that FGFR3-Abs treated animals exhibited a significantly greater cumulative pain response compared to controls (**Figure 3C**). These findings demonstrate that beyond their role as a biomarker, FGFR3-Abs induce pain behaviors in vivo.

**Figure 3.**
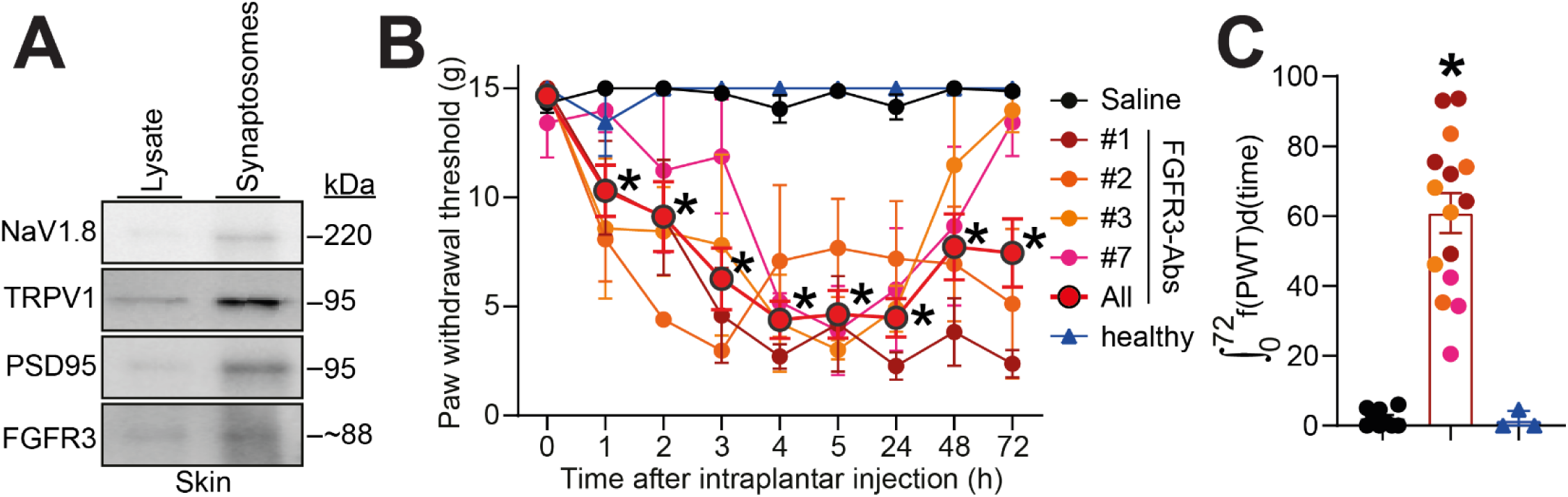
FGFR3-Abs induces mechanical hypersensitivity in vivo. (**A**) Immunoblots of skin synaptosomal fraction showing FGFR3 protein expression in skin nerve terminals. PSD95 is a marker of synapses, NaV1.8 and TRPV1 are markers of nociceptive nerve terminals. (**B**) Graph showing the paw withdrawal thresholds (PWT) of rats injected with the indicated FGFR3-Abs sera (50 µL, diluted 1:10) for up to 72h after intraplantar administration. Controls were saline and serum from a healthy control. The red symbols indicate the summary data of all FGFR3-Abs sera. Error bars indicate mean ± SEM, *p<0.05, two-way ANOVA, n= 3-6 rats per group (**C**) Bar graph with scatter plot showing the integrated area under the curve for the data in **B**, *p<0.05, Mann-Whitney test.

### FGFR3 Autoantibodies Do Not Alter Baseline Sensory Neuron Responsiveness

To investigate the mechanisms by which FGFR3-Abs could sensitize mechanical thresholds in vivo, we performed live-cell calcium imaging on cultured rat DRG neurons[22] following a 30 min exposure to 1/100 dilution of FGFR3-Abs-positive sera. Sensory neuron function was tested using depolarizing and receptor-specific stimuli, including high-potassium solution (90 mM KCl, voltage gated calcium channels), ATP (10 µM, P2X and P2Y channels), capsaicin (100 nM, TRPV1), and hypoosmotic challenge (170 mOsm, stretch gated channels). Neurons responded robustly to 90 mM KCl-induced depolarization, with no significant differences in peak calcium responses between FGFR3-Abs treated and control groups (**Figures S3A–B**). Similarly, neuronal activation by ATP (**Figure S3C–D**), capsaicin (**Figure S3E–F**), and a stretch inducing hypoosmotic challenge (**Figure S3G–H**) was comparable across conditions, indicating that FGFR3-Abs do not acutely enhance or suppress baseline sensory neuron responsiveness. These results suggest that FGFR3-Abs do not alter sensory neuron response to environmental stimuli and calcium responses under resting conditions.

### FGFR3 Autoantibodies Increase Sensory Neuron Excitability

Since FGFR3-Abs induce mechanical hypersensitivity in vivo, we next investigated whether they directly alter sensory neuron excitability, a hallmark of pain sensitization in both rodents and humans[23]. Whole-cell patch-clamp recordings were performed on DRG neurons treated with FGFR3-Abs-positive sera from two different patients (#7 and #10, 1/100 dilution for 30 min prior to recording) or a control serum obtained from the blood bank labeled as healthy control (**Figure 4A**). Neurons exposed to FGFR3-Abs-positive sera consistently exhibited increased action potential firing in response to incremental current injections compared to controls (**Figure 4A–B**). Both tested patient sera (#7 and #10) significantly enhanced neuronal excitability compared to neurons treated with PBS. Neurons treated with the serum from a healthy control showed no significant change in action potential firing. Rheobase, the minimal current required to elicit an action potential, was significantly reduced in FGFR3-Abs-treated neurons compared to controls (**Figure 4C–D**), further supporting an increase in excitability. Resting membrane potential remained unchanged across all conditions (**Figure 4E**), indicating that the observed hyperexcitability was not due to alterations in baseline membrane properties. These findings suggest that FGFR3-Abs enhance sensory neuron excitability by lowering the threshold for action potential generation, rheobase, providing a potential mechanistic link between FGFR3-Abs and pain hypersensitivity.

**Figure 4.**
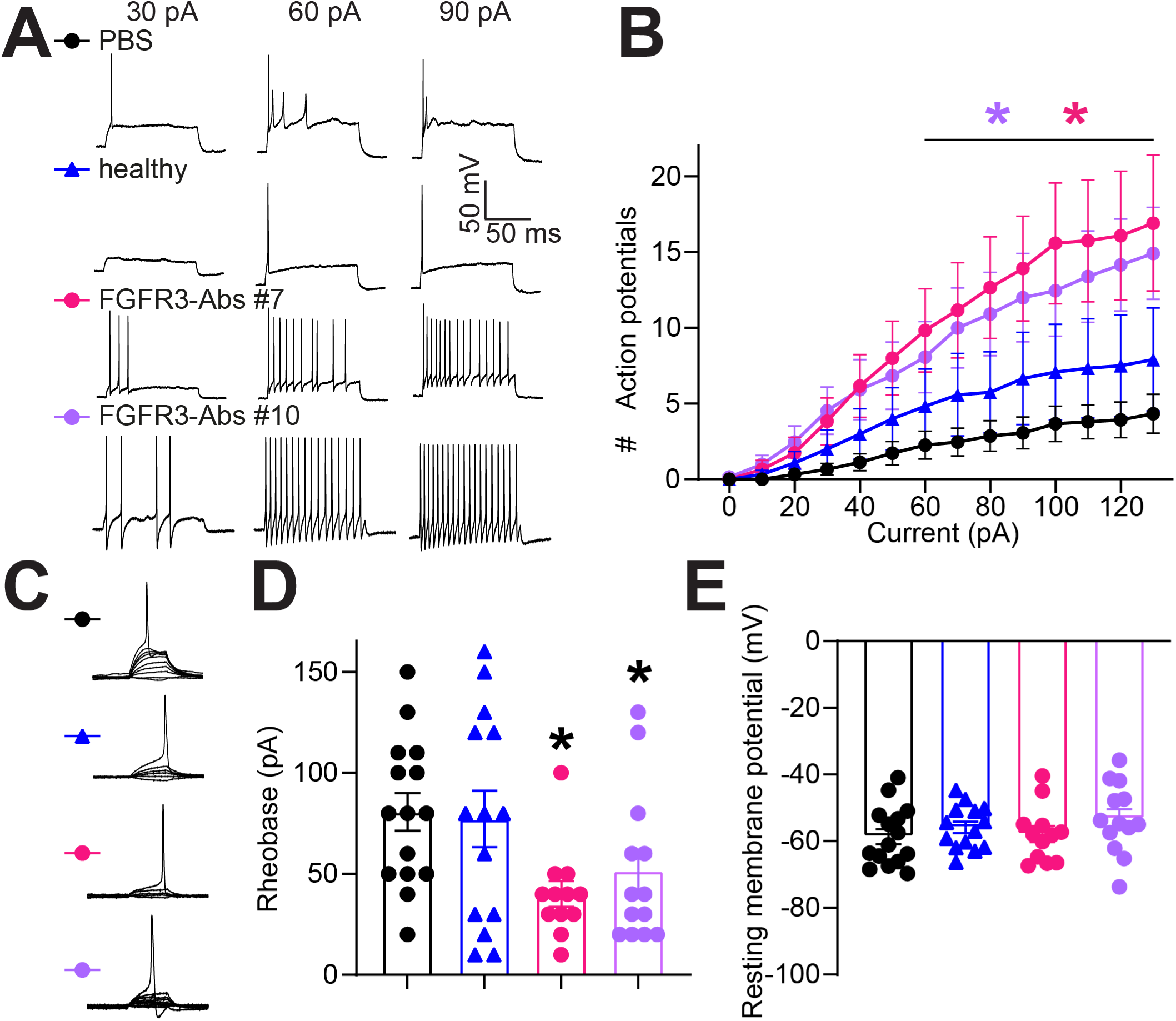
FGFR3-Abs induces hyperexcitability of rat sensory neurons. (**A**) Representative recordings of evoked action potentials at 30, 60, and 90 pA current steps from small-diameter dorsal root ganglia (DRG) neurons treated with the indicated serum (1/100 dilution) for 30 min. (**B**) Graph showing the number of evoked action potentials in response to 0-100 pA of injected current recorded from rat DRG neurons treated with either PBS as a control, FGFR3-Abs, or healthy control sera. Error bars indicate mean ± SEM, *p<0.05, Mann-Whitney test, n=12-15 cells per group. (**C**) Traces and (**D**) bar graph with scatter plot showing decreased rheobase in neurons treated with FGFR3-Abs sera compared to control. Rheobase was unchanged in neurons treated with serum from a healthy control. Error bars indicate mean ± SEM, *p<0.05, Mann-Whitney test, n=12-15 cells per group. (**E**) Bar graph with scatter plot of the resting membrane potential of neurons across all experimental conditions. Error bars indicate mean ± SEM, n=12-15 cells per group.

### FGFR3 Autoantibodies Activate MAPK Signaling in DRG

Thus far, we found that FGFR3-Abs elicit sensory neuron hyperexcitability and mechanical hypersensitivity in rats. FGFR3 signaling pathway is described to involve the MAPK p38, ERK, and c-Jun N-terminal kinase (JNK)[6]. These signaling pathways are common drivers of chronic pain and sensitize DRG neuron function[24]. To determine whether FGFR3-Abs engage these intracellular signaling pathways, we analyzed the level of activated (phosphorylated) MAPK in DRG from rats injected with FGFR3-Abs-positive sera in the paw and collected 2 hours after injection when mechanical hypersensitivity reaches a peak (**Figure 3**). As control we used the same serum after immunoglobulin depletion via adsorption on protein G beads (**Figure S4A**). Injection of FGFR3-Abs-containing serum led to a significant increase in phosphorylated ERK (p-ERK), phosphorylated JNK (p-JNK), and phosphorylated p38 (p-p38) compared to IgG-depleted serum controls (**Figure S4A–B**). Total levels of ERK, JNK, and p38 remained unchanged, indicating that FGFR3-Abs specifically enhanced MAPK phosphorylation (activation) rather than altering overall protein expression. These results suggest that FGFR3-Abs promote activation of MAPK pathways in DRG neurons in vivo, providing a mechanistic link between autoantibody exposure and enhanced neuronal excitability.

### FGFR3 Gene Editing with CRISPR Prevents Autoantibody-Induced Hyperexcitability and Hypersensitivity

We demonstrated that FGFR3-Abs induce mechanical hypersensitivity (**Figure 3**) and sensory neuron hyperexcitability (**Figure 4**). However, this does not elucidate whether autoantibodies act via binding to FGFR3 or via other membrane proteins such as Fc receptors[25]. To determine whether FGFR3 is required for the increased excitability induced by FGFR3 autoantibodies (**Figure 4**), we used CRISPR-Cas9 to genetically delete FGFR3 in sensory neurons and assessed their electrophysiological properties following exposure to FGFR3-Abs from patient #1. The CRISPR constructs were packaged into lentiviruses and added to cultured DRG neurons. We validated with both a commercial FGFR3 autoantibody and human FGFR3-Abs serum that FGFR3 was efficiently lost 48 hours after lentiviral infection (**Figure 5A-B**). Patch-clamp recordings revealed that FGFR3 knockout (FGFR3 CRISPR) neurons exhibited action potential firing properties comparable to control neurons treated with PBS, showing that FGFR3 expression does not alter basal firing properties (**Figure 5C-D**). However, FGFR3 CRISPR treated neurons failed to display increased excitability after exposure to FGFR3-Abs (**Figure 5C-D**). Rheobase was unchanged between control and FGFR3 CRISPR neurons (**Figure 5E-F**). These results demonstrate that FGFR3 deletion selectively abolished FGFR3-Abs induced hyperexcitability without altering baseline membrane properties and that FGFR3 expression is necessary for the increased sensory neuron excitability by FGFR3-Abs. Next we administered the FGFR3 CRISPR directly into the sciatic nerve of rats to selectively and unilaterally edit FGFR3 in DRG neurons. Rats were then injected intraplantarly with FGFR3-Abs patient sera as before (**Figure 3**) and mechanical thresholds measured. FGFR3 gene editing had no impact on basal mechanical thresholds, showing that FGFR3 expression in DRG neurons does not actively regulate sensitivity to an innocuous mechanical stimulus. As before (**Figure 3**), FGFR3 serum injection induced mechanical hypersensitivity on animals that received a control CRISPR injection (**Figure 5G-H**). FGFR3 gene editing prevented autoantibody induced mechanical hypersensitivity. This reinforces our observations that FGFR3-Abs act via binding to the FGFR3 receptor to sensitize mechanical thresholds. Altogether, our results suggest that FGFR3 signaling is a key driver of neuronal hyperactivity in response to autoantibody exposure, reinforcing its role in sensory hypersensitivity.

**Figure 5.**
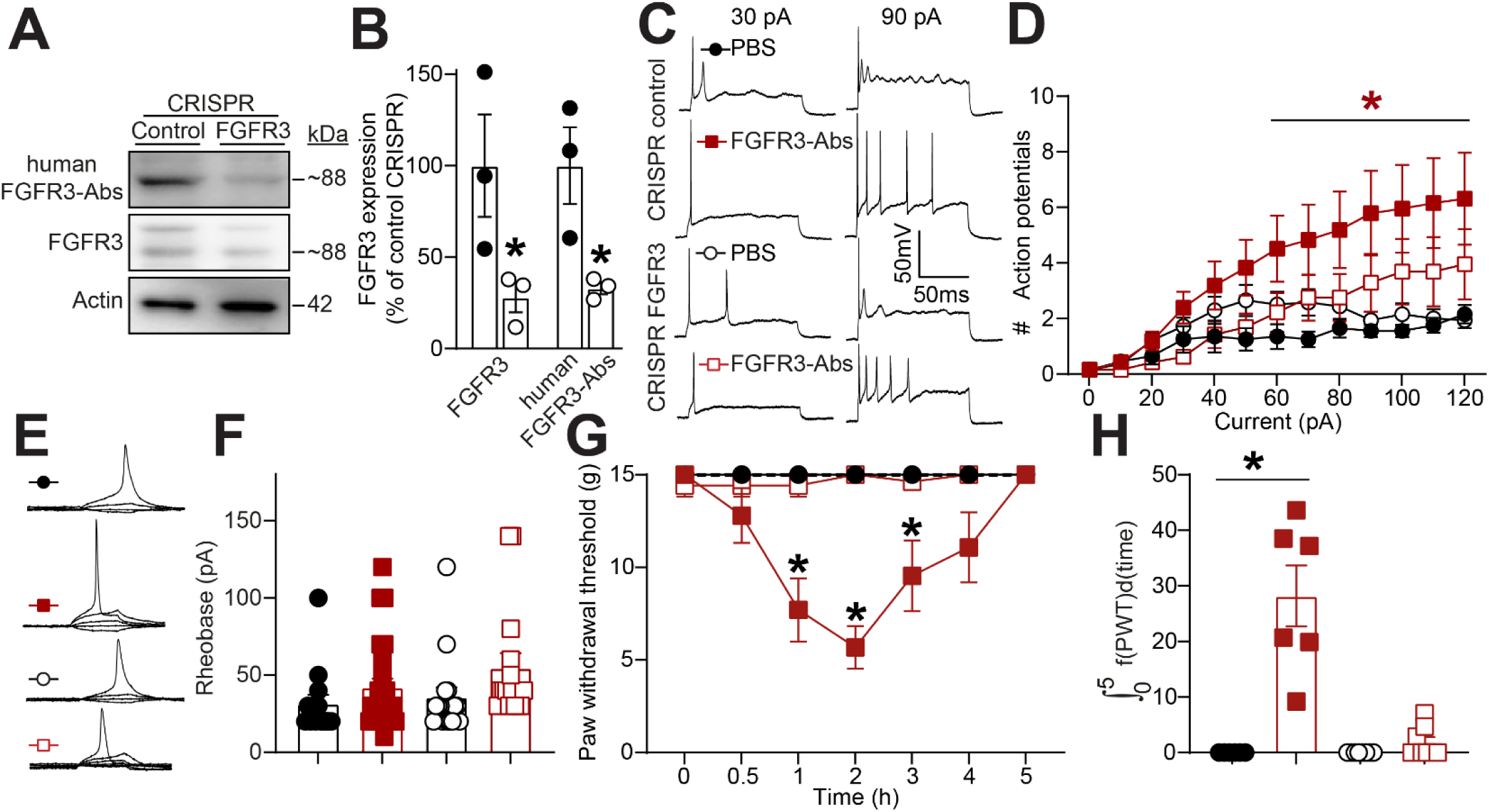
CRISPR mediated gene editing of FGFR3 prevents FGFR3-Abs driven neuronal hyperexcitability. (**A**) Immunoblots of FGFR3 expression detected with a commercial antibody or human FGFR3-Abs in cultured DRG neurons infected with CRISPR control or targeting FGFR3 as indicated. (**B**) Bar graph with scatter plot of FGFR3 expression level detected with indicated antibodies in cultured DRG neurons after FGFR3 gene editing. Error bars indicate mean ± SEM, *p<0.05 Student’s t-test, n= 3 wells per group. (**C**) Representative traces from DRG neurons with PBS, FGFR3 CRISPR alone or in conjunction with FGFR3-Abs (#1 and #3 combined). (**D**) Quantification of the number evoked action potentials in response to 0-130 pA of injected current. Error bars indicate mean ± SEM, Mann-Whitney test, n=12-17 cells per group. (**E**) Traces and (**F**) summary of rheobase measurements in all experimental conditions. Error bars indicate mean ± SEM, *p<0.05 Kruskal-Wallis test, n=12-17 cells per group. (**G**) Graph showing the paw withdrawal thresholds (PWT) of rats injected intraneurally with either control or FGFR3 CRISPR and then with FGFR3-Abs (#1 and #3 combined) (50 µL, diluted 1:10). Error bars indicate mean ± SEM, *p<0.05, two-way ANOVA, n= 6-8 rats per group (**H**) Bar graph with scatter plot showing the integrated area under the curve for the data in **G**, *p<0.05, Mann-Whitney test.

### FGFR3 Autoantibodies Target Specific Extracellular and Intracellular Epitopes

Next we wanted to address that FGFR3-Abs act via their antigen binding fragment (Fab) to sensitize DRG neurons and pain responses. We synthesized a peptide array containing 15-mer peptides spanning the entire human FGFR3 sequence in 3 amino-acid increments (**Figure 6A**). Peptide arrays were hybridized with patient sera and signal developed with an anti-human fluorescent secondary antibody. This let us detect binding domains located mainly on the extracellular domain (ECR) with additional sequences located in the juxtamembrane domain (JMD) and slightly overlapping with the tyrosine kinase domain (**Figure 6A-B**). These results are consistent with a previously published epitope mapping of FGFR3-Abs[9]. However, we found a consistent and strong signal targeting the ECR domain suggesting that FGFR3-Abs could predominantly bind to the ligand binding portion of FGFR3 to elicit signaling. To further investigate autoantibody binding to FGFR3 in sensory neurons, we performed immunolabeling on PFA-fixed DRG neurons in the absence or presence of the detergent Triton X-100, which permeabilizes the plasma membrane. In non-permeabilized conditions (-T-X100), FGFR3-Abs from two different patients labeled the neuronal surface, indicating recognition of extracellular epitopes (**Figure 6C, see inset**). In permeabilized conditions (+ T-X100), intracellular FGFR3 staining was also detected, confirming additional autoantibody interactions with cytoplasmic regions (**Figure 6C**). Quantification of fluorescence intensity revealed significant FGFR3-Abs binding in both non-permeabilized and permeabilized conditions (**Figure 6D–E**). These results demonstrate that FGFR3-Abs recognize extracellular epitopes on FGFR3. This observation infers that FGFR3-Abs could directly sensitize DRG neurons and pain via binding to the ECR of FGFR3.

**Figure 6.**
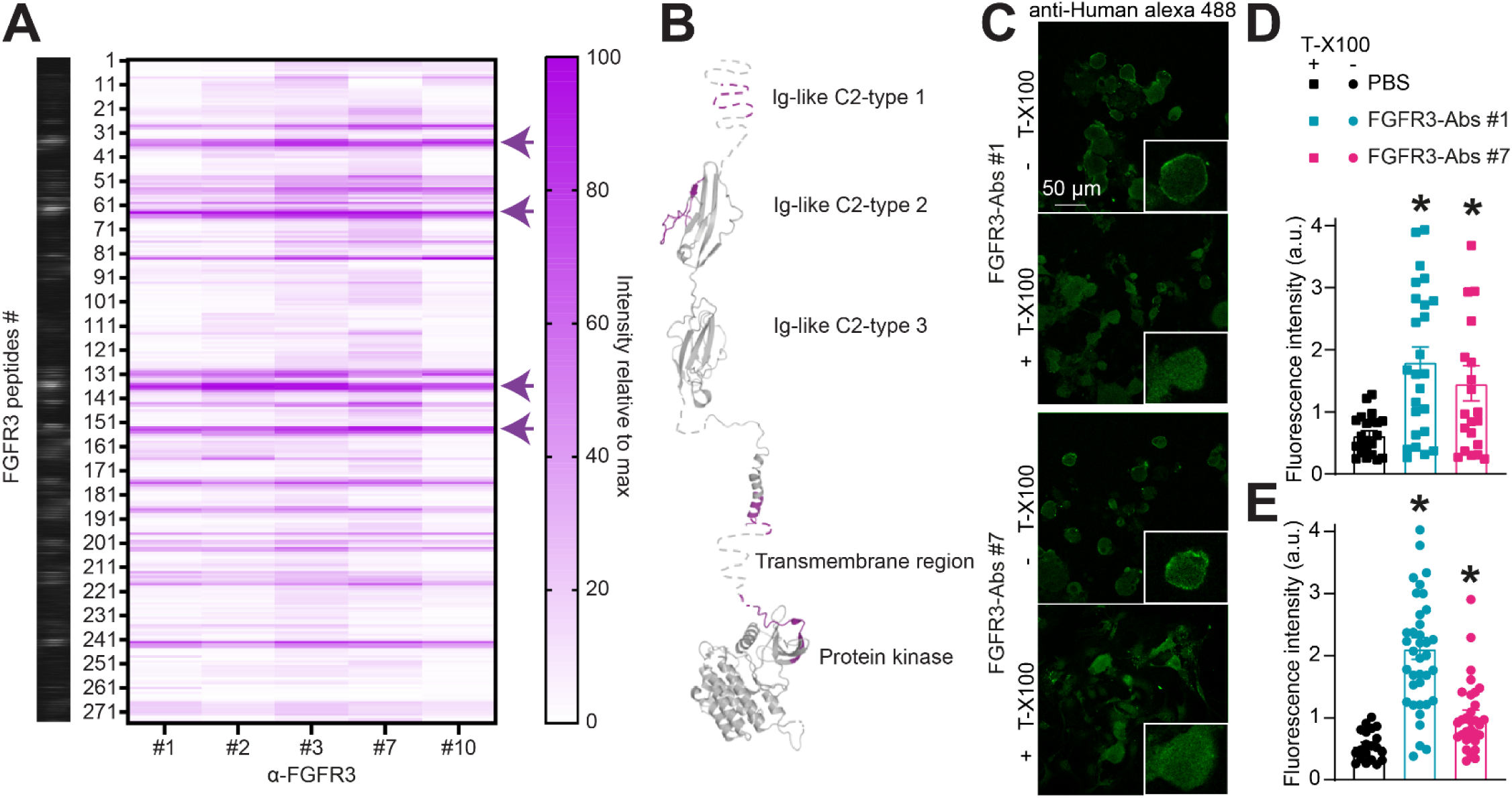
FGFR3-Abs bind to FGFR3 extracellular domain. (**A**) Heatmap of the immunoreactive hotspots detected by human FGFR3-Abs-positive sera hybridized to 15-mer segments of human FGFR3 (UniProt ID: P22607). Four main epitopes were identified within the extracellular region of FGFR3 indicated by arrows. (**B**) Spatial representation of the FGFR3 protein domains and epitopes identified in **A,** mapped on mapped on representative structures PDBID: 3GRW [40], PDBID:2LZL [41] and PDBID: 6LVL [42]. (**C**) Representative immunocytofluorescent images of dissociated DRG neurons immunolabeled with human FGFR3-Abs sera. Different permeabilization conditions (with or without triton X-100) were used to limit the access of autoantibodies to the extracellular or intracellular space. Scale bar is 50 µm. Bar graph with scatter plot of the FGFR3-Abs fluorescence intensity measured in (**D**) permeabilized or (**E**) non-permeabilized sensory neurons. Error bars indicate mean ± SEM, *p<0.05 Kruskal-Wallis test, n=20-38 cells per group.

### FGFR3 Autoantibody-Induced Hyperexcitability is Mediated by Extracellular Epitope Binding

Given that FGFR3-Abs recognize both extracellular and intracellular domains, we tested whether blocking extracellular epitope binding could prevent autoantibody-induced sensory neuron hyperexcitability. To test this, we generated four peptides based on the four ECR epitope sequences to block the Fab portion of autoantibodies. DRG neurons were treated for 30 min with either PBS, FGFR3-Abs #1 (1/100 dilution), ECR peptides (100 ng/ml) or with a mixture of FGFR3-Abs #1 and ECR blocking peptides and whole-cell patch-clamp recordings were performed. Consistently with our other data (**Figure 4**), neurons treated with FGFR3-Abs #1 alone exhibited a robust increase in action potential firing in response to incremental current injections compared to the control (**Figure 7A–B**). However, while the ECR peptides had no significant action by themselves, peptides curbed the increased excitability due to FGFR3-Abs #1 (**Figure 7A-B**), restoring firing frequency to near-control levels. Rheobase measurements again showed that FGFR3-Abs #1 significantly lowered the current threshold required for action potential generation, which was not impacted by ECR blocking peptides (**Figure 7C–D**). Resting membrane potential remained unchanged across conditions (**Figure 7E**). Similarly, blocking FGFR3-Abs with ECR peptides prevented the development of autoantibody induced mechanical hypersensitivity (**Figure 7G-H**). We additionally tested if peptides based on the JMD [9] could prevent sensory neuron hyperexcitability induced by FGFR3-Abs #1. JMD peptides alone had the surprising action of increasing excitability compared to baseline (**Figure S5A-D**). The co-treatment with FGFR3-Abs #1 was not distinguishable from JMD peptides alone (**Figure S5A-D**). In vivo, JMD peptides had no impact on FGFR3-Abs induced mechanical hypersensitivity (**Figure S5E-F**). Altogether, these results indicate that FGFR3-Abs-induced neuronal hyperexcitability and hypersensitivity is primarily mediated through extracellular epitope binding. Blocking these interactions with FGFR3 ECR peptides prevented autoantibody induced sensitization, suggesting a potential therapeutic strategy for mitigating autoantibody-induced sensory dysfunction.

**Figure 7.**
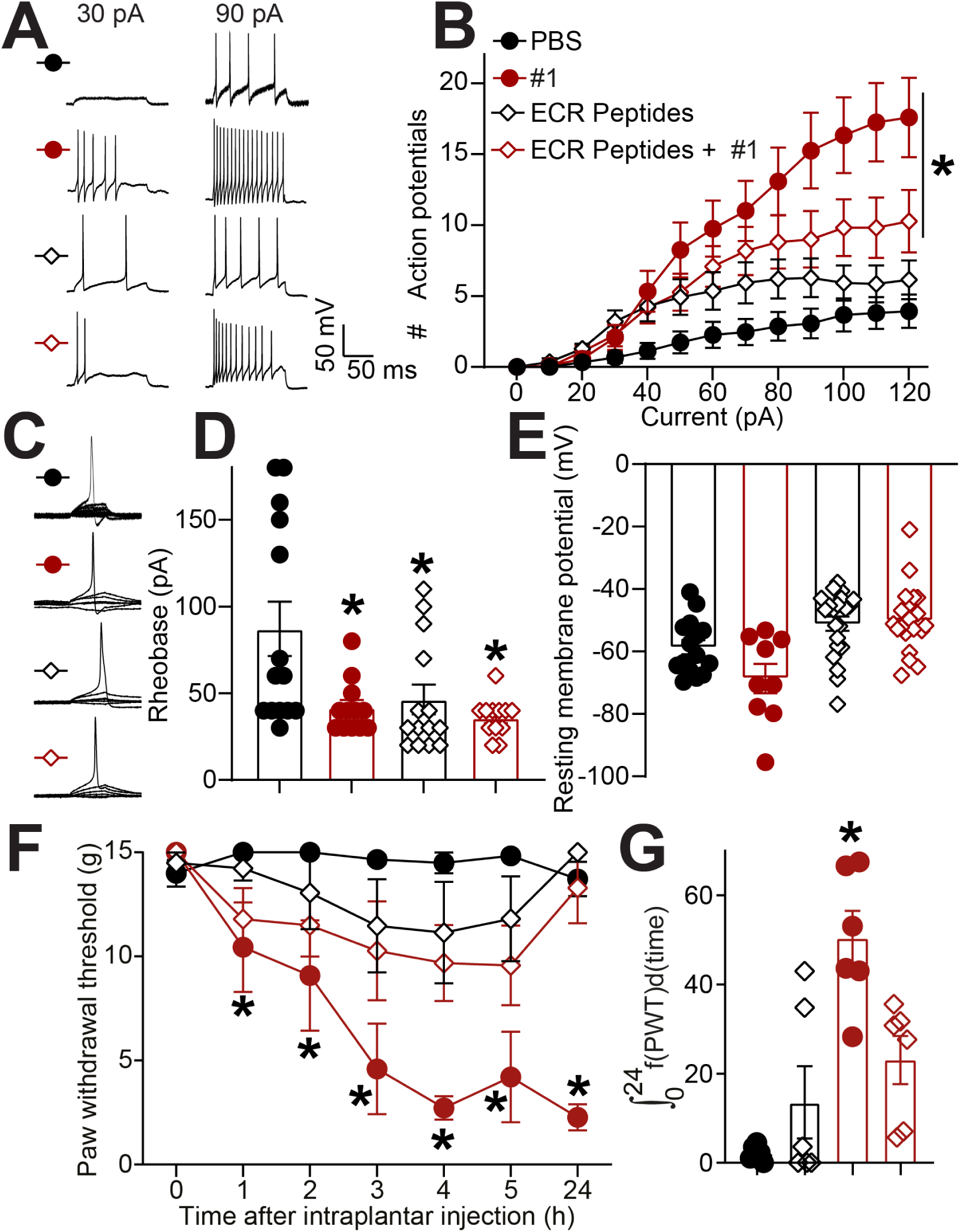
ECR-binding peptides block neuronal hyperexcitability induced by FGFR3-Abs in rat sensory neurons. (**A**) Representative recordings of evoked action potentials recorded from small diameter DRG neurons treated with serum from FGFR3-Abs-positive patient #1 (1/100 dilution) in combination with or without 100 ng/ml of ECR peptides as indicated, for 30 min in response to depolarizing current injection of 30 and 90 pA. (**B**) Quantification of the number evoked action potentials in response to 0-120 pA of injected current. Mean ± SEM, n=12-21 cells per condition, *p<0.05, Mann-Whitney test. (**C**) Traces and (**D**) bar graph with scatter plot showing decreased rheobase in cells treated with the serum from FGFR3-Abs #1 and/or ECR peptides as indicated. Mean ± SEM, n=12-21 cells per condition, *p<0.05, Kruskal-Wallis test. (**E**) Bar graph with scatter plot of the resting membrane potential of neurons from each treatment group. Error bars indicate mean ± SEM, *p<0.05, Kruskal-Wallis test, n=12-21 cells per condition. (**F**) Graph showing the paw withdrawal thresholds (PWT) of rats injected with FGFR3-Abs #1 sera (50 µL, diluted 1:10) and ECR peptides (100ng/ml each) as indicated. Error bars indicate mean ± SEM, *p<0.05, two-way ANOVA, n= 6 rats per group (**G**) Bar graph with scatter plot showing the integrated area under the curve for the data in **F**, *p<0.05, Mann-Whitney test.

## Discussion

In this study, we uncover a pathogenic role for FGFR3-Abs from patients with peripheral neuropathies in the sensitization of DRG sensory neurons by provoking hyperexcitability and pain behaviors in rats. Building on clinical observations, we first noted that all FGFR3-Abs patients reported neuropathic pain and exhibited atypical features including small-fiber involvement and non-length-dependent fiber loss raising the possibility that the dorsal root ganglia are a primary site of injury. In a first step, we validated two important postulates: (*i*) FGFR3 is expressed in human DRG neurons at the transcript and protein levels, and *(ii*) FGFR3-Abs can find their target in sensory neurons. In vivo studies showed that serum from patients with FGFR3-Abs induced mechanical hypersensitivity while the serum from a healthy control had no effect. This effect on pain perception coincided with the sensitization of action potential firing in sensory neurons by FGFR3-Abs which required FGFR3 expression. Our next avenue of investigation identified extracellular and intracellular epitopes targeted by FGFR3-Abs. Peptides designed on the epitope sequences were then used to confirm that FGFR3-Abs binding to the extracellular domain of FGFR3 triggered sensory neuron hyperexcitability and mechanical hypersensitivity. Ultimately, our data unveils the role of FGFR3-Abs in sensitizing pain via binding to the extracellular domain of FGFR3 on sensory neurons leading to an activation of FGFR3 signaling. These discoveries enhance our understanding of how pain responses are sensitized in patients with SNN and SFN by auto-immunoreactivity of FGFR3-Abs against sensory neurons consequently lowering the threshold for nociceptive action potential firing.

Our endeavors build on the discovery of autoantibodies against FGFR3 as a biological marker in patients with sensory neuronopathy and small-fiber neuropathy[3a] with a high prevalence of pain[4]. With our results showing that FGFR3-Abs directly sensitize DRG neurons and pain behaviors in vivo, we can conclude that these autoantibodies, beyond their role as biomarkers, might functionally promote pain[26]. These observations are in line with recent clinical studies that failed to detect clear beneficial outcomes for patients with FGFR3-Abs-positive SFN treated with intravenous immunoglobulin (IVIg)[27]. IVIg functions by saturating Fc receptors to prevent the activation of immune cells and the complement cascade[28]. The lack of apparent efficacy of IVIg treatment suggests that symptoms characterizing FGFR3-Abs-positive neuropathies are independent of Fc-mediated effects. From our characterization of FGFR3-Abs epitopes, we identified blocking peptides capable of saturating the Fab region of autoantibodies, which prevented FGFR3-Abs-induced increases in sensory neuron excitability. Given these results, we postulate that those epitope peptides could be used for therapy by saturating FGFR3-Abs autoantibodies as a form of antigen-specific immunotherapy. Another application relies in engineering Apitopes (Antigen Processing Independent epiTOPEs) which will mimic the CD4^+^ T-cell epitopes to induce immune tolerance towards self-antigens [29]. With our approach of high density peptide mapping, personalized medicine can be achieved by facile epitope sequence identification to then inform the production of Apitopes tailored to each patients’ autoimmune response. We previously demonstrated the benefits of an anti-CD20 monoclonal antibody[30], which targets autoantibody-producing B lymphocytes[31] by reducing serum autoantibody levels and reversing mechanical hypersensitivity in a rat model of autoimmune neuropathy[21]. Consistent with this, a recent clinical study reported a significant improvement in visual analog scale (VAS) pain scores in FGFR3-Abs-positive SFN patients treated with the anti-CD20 monoclonal antibody Rituximab[32]. Taken together, our findings and recent clinical evidence support a shift in treatment strategies from saturating Fc receptors with IVIg to instead focus on reducing circulating autoantibody levels via use of anti-CD20 therapeutics.

It is widely accepted that autoantibodies can complex to their target, however whether autoantigen recognition by said target either is agonistic or antagonistic of the target protein is often an open question. Our results indicate that FGFR3-Abs can bind to their cognate target on DRG neurons and trigger hyperexcitability. As such, we can conclude that FGFR3 is (*i*) expressed in both rat and human DRG neurons, (*ii*) coupled to intracellular signaling pathways via ERK, p38, and JNK, (*iii*) activated by binding of FGFR3-Abs on its extracellular domain, (*iv*) involved in regulating action potential firing in sensory neurons, and (*v*) a key mediator of FGFR3-Abs induced pain behaviors. p38[33] and ERK[34] signaling pathways are directly related to increased action potential firing in chronic neuropathic pain[24, 35]. This could be linked to increased voltage gated sodium channel Nav1.8 current density via p38 mediated phosphorylation[33a]. Another contributor might be phosphorylation of the threshold channel Nav1.7 via ERK or p38, leading to facilitated sensory neuron firing[34a, 36]. Since FGFR3-Abs activate both kinases in DRG, it is likely that Nav1.7 sensitization contributes to the decreased rheobase in sensory neurons treated with human autoantibodies while increased Nav1.8 current density via p38 further enhances neuronal excitability.

Previously, FGFR3 was found in spinal astrocytes where it contributed to TNF-α expression[37]. FGFR3 expression was increased in the spinal dorsal horn of rats[37] with the spared nerve injury (SNI) model of chronic neuropathic pain[38]. However, the functional relevance of FGFR3 in this context remains uncertain as only a high dose (10 µM, intrathecal) of PD173074, an selective FGFR3 antagonist (Ic50 = 5 nM) with known off-target effects on FGFR1 (Ic50 = 21.5 nM), VEGFR2 (Ic50 = 100 nM), PDGFR (Ic50 = 17.6 µM) and c-Src (Ic50 = 19.8 µM)[39], was able to reverse mechanical hypersensitivity in SNI rats[37]. This raises concerns about whether FGFR3 or broader inhibition of receptor tyrosine kinases resulted in analgesia in this study. No reports to date have detected FGFR3-Abs in the cerebrospinal fluid (CSF) of patients[3a, 4, 9], suggesting that direct central effects of FGFR3-Abs are unlikely. In addition to astrocytes, FGFR3 was described to be expressed in other non-neuronal cell types within the DRG, including Schwann cells, satellite glia, and macrophages[5]. Given that FGFR3-Abs bind DRG neurons (**Figures 1, 6** and [3a]), our study focused on their peripheral actions on primary afferents rather than central mechanisms. Importantly, our findings indicate that FGFR3 expression in DRG neurons plays a direct role in sensory neuron hyperexcitability, suggesting that FGFR3 may contribute to heightened pain sensitivity in non-autoimmune neuropathic pain conditions as well. This distinction is crucial, as it shifts the focus from central FGFR3 signaling in astrocytes to a more functionally relevant role in sensory neurons, where FGFR3-Abs binding actively drives pain hypersensitivity. While we cannot exclude the possibility that FGFR3-Abs may influence other cells of the peripheral nervous system, our CRISPR and blocking peptide experiments demonstrate that the involvement of sensory neuron FGFR3 is sufficient to account for the autoantibody induced hypersensitivity.

A key limitation of our study was the restricted availability of FGFR3-Abs-positive sera, which prevented a comprehensive analysis for all measures across a larger cohort. However, our clinical data (n = 63) showing consistent neuropathic pain symptoms and a distinct electrophysiological profile with frequent small-fiber involvement, provides a broader context to our mechanistic findings. Additionally, it remains unclear whether our sample set reflects any sex-based bias, though no reports have indicated a sex-related prevalence of FGFR3 autoantibodies. Despite these constraints, our epitope mapping consistently identified comparable epitopic patterns for FGFR3-Abs binding to the extracellular domain of FGFR3. This together with our experiments demonstrating that blocking autoantibody binding to the extracellular domain of FGFR3 prevented pain and hyperexcitability, suggests that the detection of FGFR3-Abs against extracellular epitopes could be a predictor of painful SFN in patients. Finally, in line with prior studies that could not correlate FGFR3-Abs levels with symptom severity in SFN or SNN patients, we chose not to account for absolute autoantibody titers in our experiments. Importantly, the combination of autoantibody depletion, CRISPR editing of FGFR3, epitope-blocking experiments, and validation with a healthy control corroborate that sensory neuron hyperexcitability is driven by FGFR3-Abs binding rather than other serum components.

Our findings establish FGFR3-Abs as direct contributors to sensory neuron hyperexcitability and pain, shifting their role from mere biomarkers to active disease drivers in autoimmune sensory neuropathies. By demonstrating that epitope saturation of FGFR3-Abs prevents mechanical hypersensitivity, we suggest B-cell-targeted therapies, such as anti-CD20 monoclonal antibodies, as a promising avenue to be evaluated for treating FGFR3-Abs linked neuropathies. Moreover, the ability of FGFR3-Abs to activate FGFR3 signaling and sensitize nociceptors suggests that FGFR3 itself may represent a novel therapeutic target for modulating sensory neuron excitability beyond autoimmune neuropathies.

## Data availability

The data to support the findings of this study are available from the corresponding author upon reasonable request.

## Acknowledgements

We thank Mid America transplant with Drs. Grant Kolar, Robert W. Gereau IV and Bryan Copits for their support in facilitating access to human DRG. We thank Edel Edorh, Christine Kim, Avani Mathur and Niloufarsadat Mirian for technical support for these studies.

## Supplementary material

Supplementary material is available at *Advanced Science* online.

---

**Figure S1.**
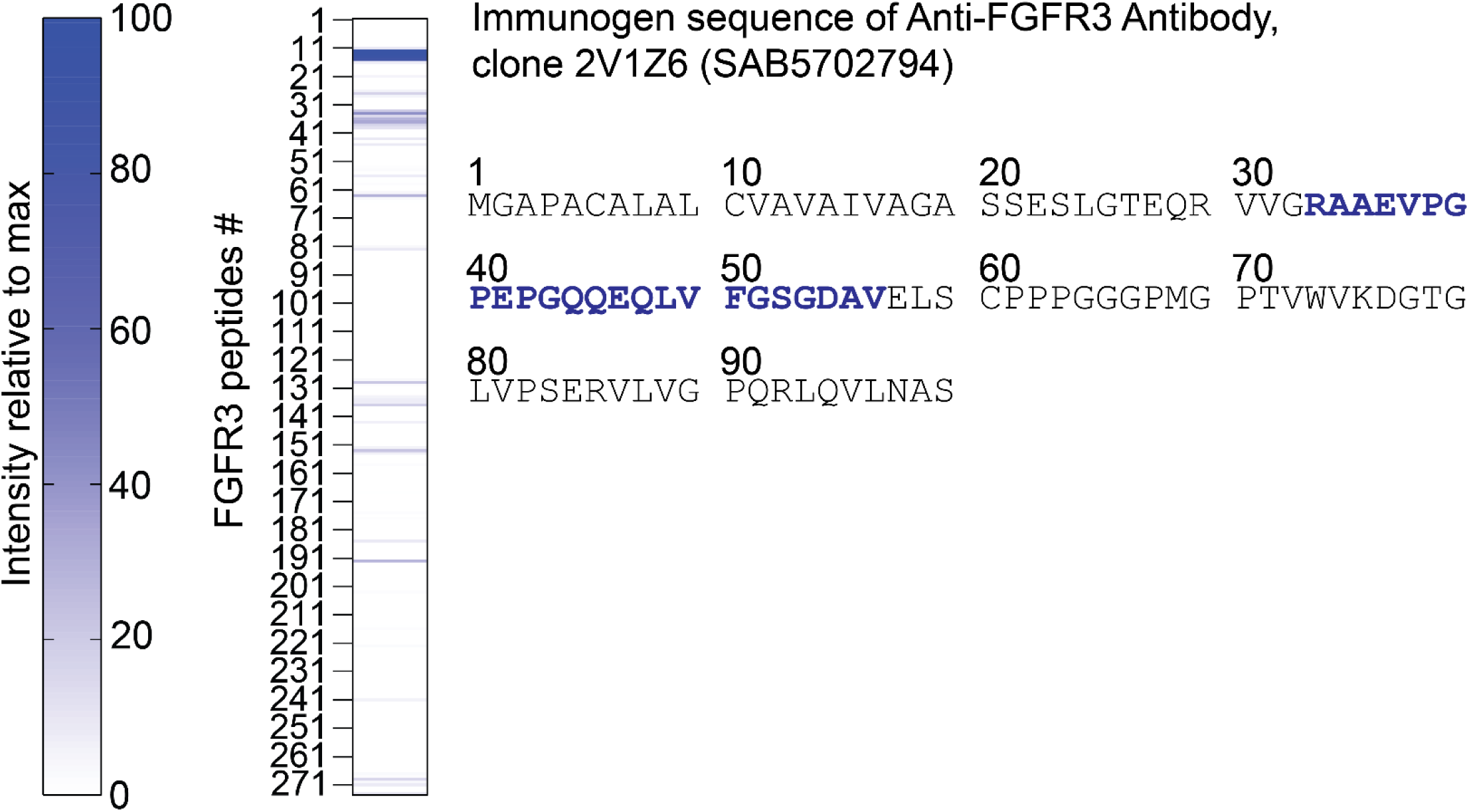
Epitope mapping of the commercial anti-FGFR3 antibody using our FGFR3 peptide array. Our peptide array spanning the N-terminal sequence of FGFR3 was probed with the commercial anti-FGFR3 antibody. The heatmap (left) shows relative signal intensity across the overlapping 15-mer peptides tiled along the human FGFR3 protein sequence. Peptides corresponding to the immunogen region (residues 30–60) exhibit the strongest binding signal. This confirms the specific reactivity of our commercial FGFR3 antibody to its intended immunogen region within the FGFR3 extracellular domain. This also confirms the validity of our peptide array strategy. Signal intensity is scaled relative to the maximum observed response.

**Figure S2.**
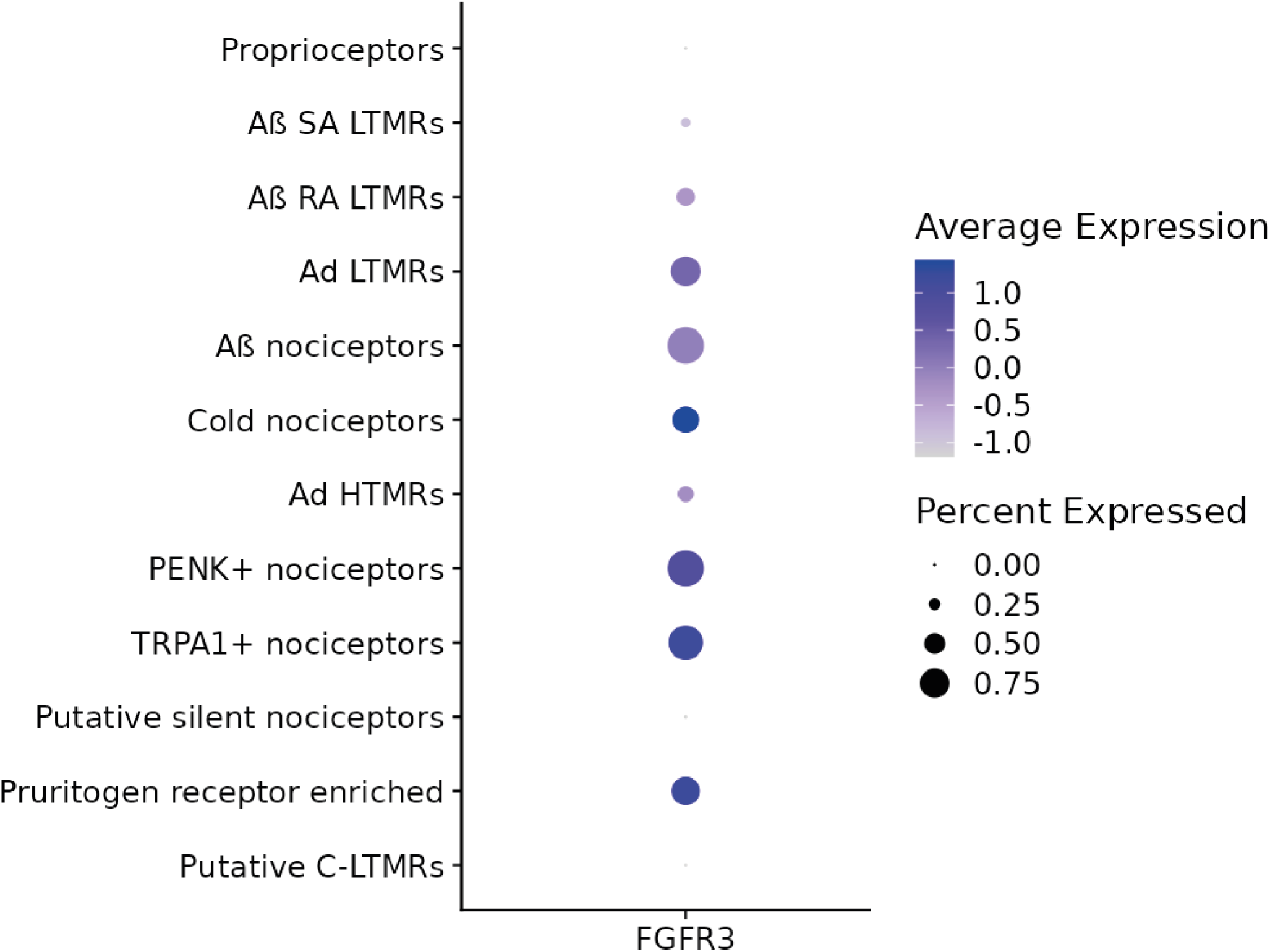
FGFR3 expression across human DRG neuronal subtypes identified via spatial transcriptomics. Bubble plot depicting the expression of FGFR3 in human dorsal root ganglion (DRG) neuronal subtypes. The size of each circle indicates the percentage of cells within a given subtype expressing FGFR3, while the color scale reflects the average expression level (blue = higher expression, lighter shades = lower expression). Data were mined from [19].

**Figure S3.**
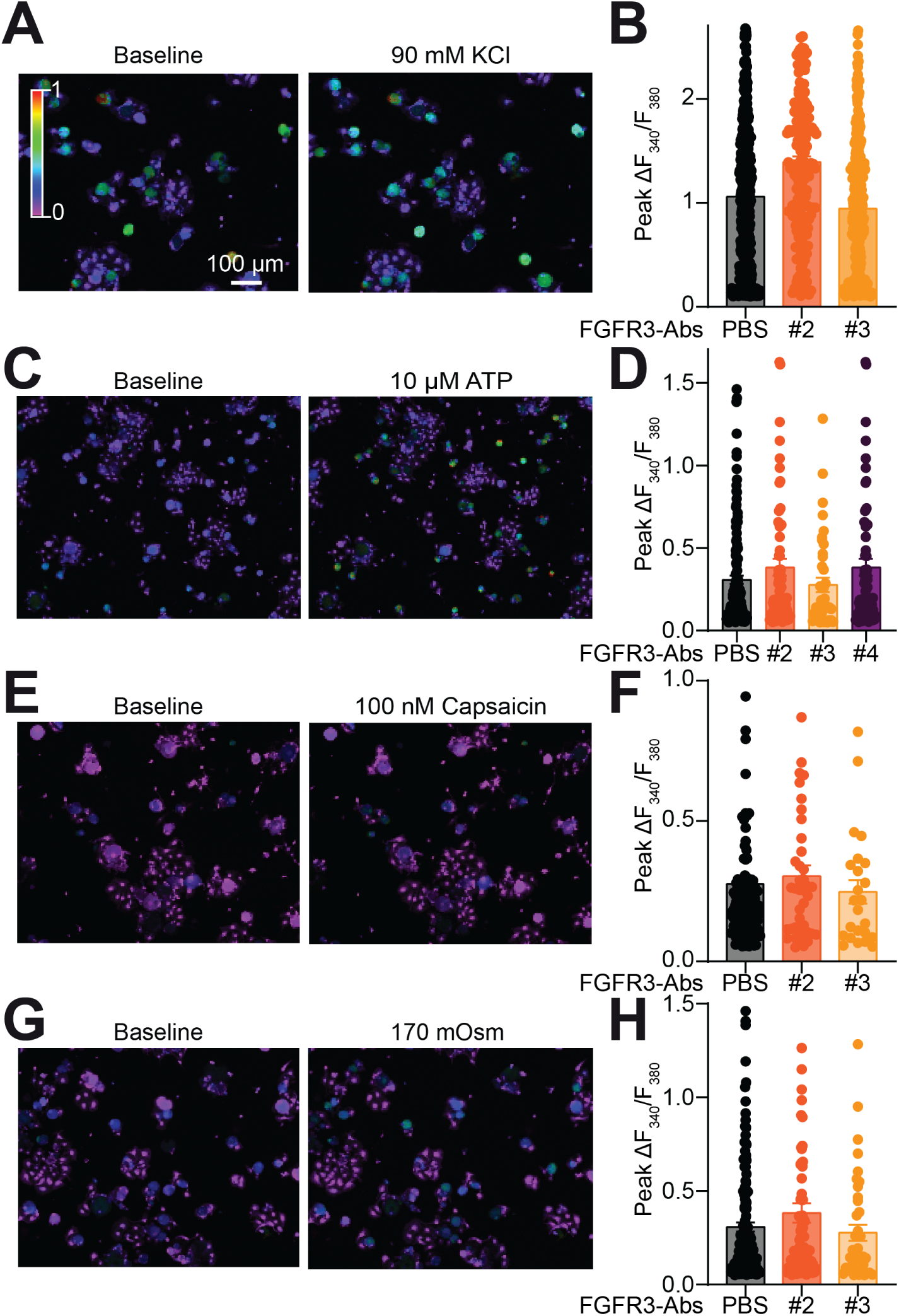
Human FGFR3-Abs autoantibodies have no effect on sensory neuron functions. Rat DRG neurons loaded with the Ca^2+^ sensitive dye Fura-2AM were examined for their response to (**A-B**) depolarization (90mM KCl), (**C-D**) 10µM ATP, (**E-F**) 100nM Capsaicin, and (**G-H**) 170 mOsm (hypoosmotic induced stretch). Following incubation with FGFR3-Abs (1/100 dilution, 30 min), dissociated DRG neurons were exposed to the above triggers and Ca^2+^ influx was measured and plotted following the Fura-2AM fluorescence ratio (F_340_/F_380_). The responses of DRG neurons were simultaneously recorded in response to the respective treatments. Baseline images were taken prior to trigger application or at peak after application of indicated trigger. In all cases 90 mM KCl was used an indicator of viability and only cells responding to depolarization were used for analysis. The color scale indicates the value of the Fura-2AM fluorescence ratio (F_340_/F_380_) with red indicating the highest concentration of intracellular Ca^2+^. Mean ± SEM, *p< 0.05 compared to control, Kruskal-Wallis test.

**Figure S4.**
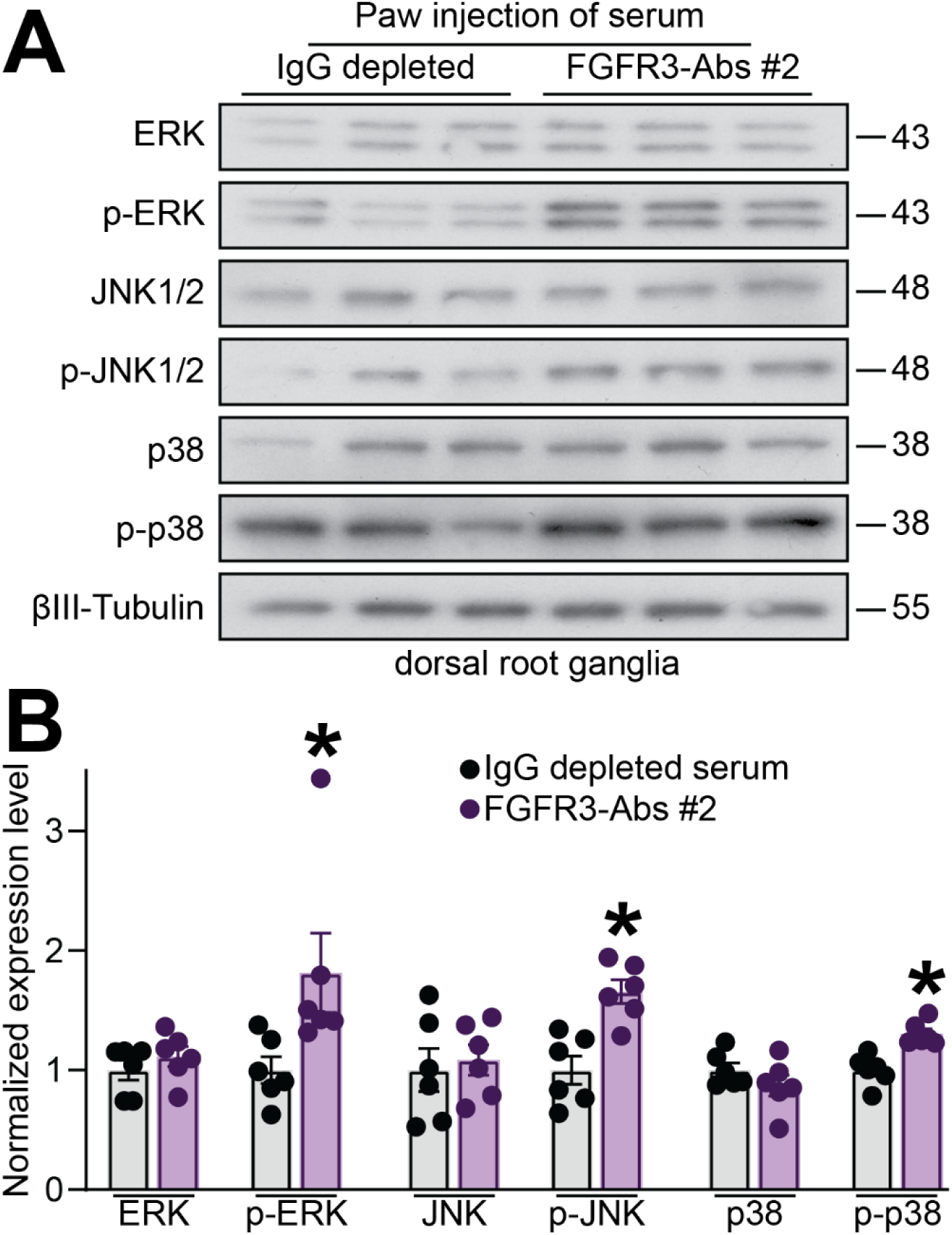
Intraplantar injection of FGFR3-Abs activated FGFR3 downstream signaling pathways. Rats were injected in the paw with FGFR3-Abs-positive serum (50 µL, diluted 1:10) or IgG depleted serum. DRG were harvested 2h after injection. (**A**) Representative immunoblots depicting levels of ERK, JNK1/2, and p-p38 as well as their respective phosphorylated counterparts. (**B**) Bar graph with scatter plot showing levels of phosphorylated ERK, JNK, and p38 in DRG from rats injected as indicated. Mean ± SEM, *p<0.05, Mann-Whitney test, n=6 rats per group.

**Figure S5.**
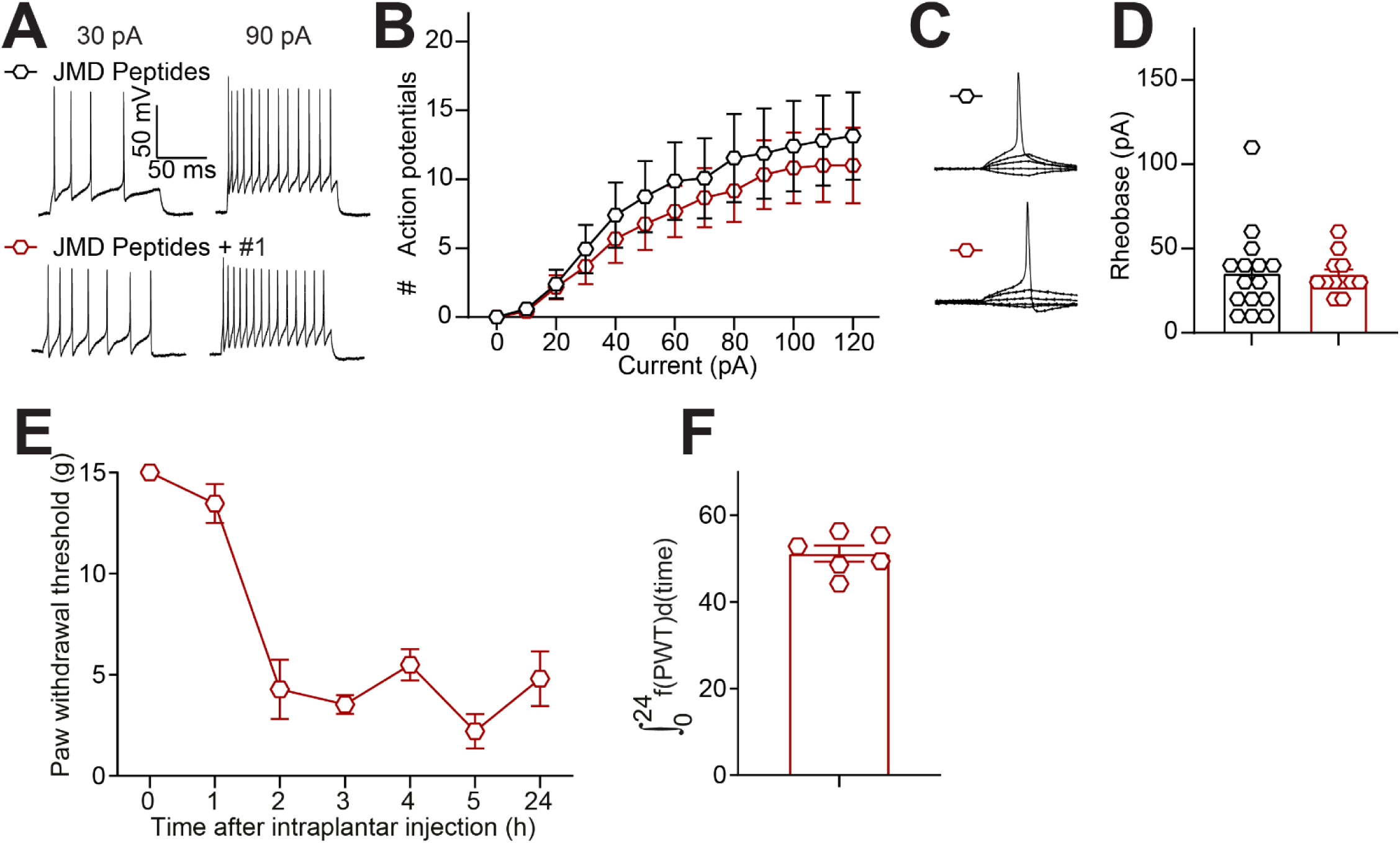
Juxtamembrane domain targeting peptides fail to prevent FGFR3-induced neuronal hyperexcitability. Treatment of DRG neurons with peptides designed on the juxtamembrane domain (JMD, 100 ng/ml) alone or in combination with FGFR3-Abs serum. (**A**) Representative traces of DRG neurons in either treatment group exhibit similar firing patterns at 30 and 90 pA current steps and (**B**) number evoked action potentials in response to 0-120 pA of injected current. (**C**) Representative traces and (**D**) Bar graph with scatter plot showing comparable rheobase between treatments. Mean ± SEM, Mann-Whitney test, n=12-15 cells per group. (**E**) Graph showing the paw withdrawal thresholds (PWT) of rats injected with FGFR3-Abs sera (50 µL, diluted 1:10) with JMD peptides (100ng/ml each). Error bars indicate mean ± SEM, n= 6 rats per group (**F**) Bar graph with scatter plot showing the integrated area under the curve for the data in **E**.

**Table S1:** Anti-FGFR3 positive sera used in this study. Sex, type of neuropathy and age at diagnostics are indicated. All patients had a type of neuropathy and those who reported pain are indicated.

| Patient # | Diagnostic | Onset | Pain | Anti-FGFR3 | Other Ab | Sex | Age |
| --- | --- | --- | --- | --- | --- | --- | --- |
| 1 | Polyneuropathy | Acute | Yes | Positive | Negative | M | 65 |
| 2 | SNN | Acute | Unknown | Positive | Negative | M | 59 |
| 3 | SFN | Subacute | Yes | Positive | Negative | F | 62 |
| 4 | Sensory neuropathy with motor disorder | Progressive | Yes | Positive | Negative | M | 84 |
| 7 | SNN | Subacute | Yes | Positive | Negative | M | 35 |
| 10 | SNN | Progressive | Yes | Positive | anti-gangliosides | F | 78 |

**Table S2:** Human DRG used in this study. Age sex and cause of death at the time of organ donation.

| Age | Sex | Cause of death | Mechanism of death |
| --- | --- | --- | --- |
| 48 | male | Stroke | Intracranial Hemorrhage |
| 25 | Female | Stroke | Intracranial Hemorrhage |
| 61 | Male | Stroke | Intracranial Hemorrhage |
| 44 | Male | Anoxia | Drug Intoxication |
| 65 | Male | Anoxia | Cardiovascular |
| 45 | Female | Stroke | Intracranial Hemorrhage |
| 65 | Female | Stroke | Intracranial Hemorrhage/stroke |
| 74 | Male | Stroke | Intracranial Hemorrhage/stroke |
| 43 | female | Stroke | Intracranial Hemorrhage/stroke |
| 45 | Male | Anoxia | Cardiovascular |
| 62 | Male | Anoxia | Cardiovascular |
| 54 | Female | Anoxia | Drug Intoxication |
| 31 | Female | Anoxia | Drug Intoxication |
| 38 | Male | Anoxia | Drug Intoxication |
| 43 | Female | CNS Tumor | Death from Natural Causes |
| 62 | Male | Stroke | Intracranial Hemorrhage/stroke |
| 39 | Male | Anoxia | Drug Intoxication |
| 43 | Female | Stroke | Intracranial Hemorrhage/stroke |
| 63 | Male | Stroke | Intracranial Hemorrhage/stroke |
| 51 | Female | Anoxia | Drug Intoxication |
| 66 | Male | Head trauma | Gunshot wound |
| 59 | male | Head trauma | Blunt Injury |
| 53 | female | Cerebrovascular/Stroke | Intracranial Hemorrhage/stroke |
| 61 | male | Stroke | Intracranial Hemorrhage/stroke |
| 19 | female | Anoxia | none of the above |
| 64 | Female | Cerebrovascular/Stroke | Intracranial Hemorrhage/stroke |
| 59 | male | Anoxia | Cardiovascular |
| 23 | male | Anoxia | intoxication |
| 44 | Female | Head trauma | Blunt Injury |

